# Dynamics of the B-Cell gene regulatory network in differentiation determine evolutionary trajectories of childhood leukaemogenesis

**DOI:** 10.64898/2026.05.21.724796

**Authors:** Matthieu Bouguéon, Jason Wray, Tariq Enver, Benjamin A Hall

## Abstract

B-cell precursor acute lymphoblastic leukaemia (BCP-ALL) is the most frequent paediatric malignancy. Despite extensive molecular and cellular characterization a mechanistic model of leukaemogenesis has not yet been developed. Here we present a multi-valued logical model of B-cell differentiation, centred on a core pentad of transcription factors whose dysregulation results in the developmental arrest characteristic of BCP-ALL.

Whilst B-cell maturation follows a fixed and sequential differentiation process, leukaemic transformation is driven by stochastic genetic insults. Integrating *BCR-ABL1* as the initiating event, we model alternative evolutionary trajectories by introducing secondary mutations at different time points within synchronous simulations. We demonstrate that the combination, order and timing of secondary mutations dictate maturation arrest points and fitness, explaining how different patterns of mutations seen in patients may arise and in turn influence disease severity. This platform provides a generalisable framework to model driver mutations in their developmental context to predict evolutionary trajectories in cancer.

## 1 Introduction

B-cell precursor acute lymphoblastic leukaemia (BCP-ALL) is the most common paediatric malignancy, representing approximately 80% of childhood leukaemia cases with peak incidence in children aged 2 to 5 years [1, 2]. Genomic studies provide compelling evidence that initiating events, frequently arise in utero during fetal haematopoiesis [3, 4]. These first-hit alterations are necessary but insufficient for overt disease, creating a persistent preleukaemic clone; transformation typically requires secondary genetic alterations.

BCP-ALL is thought to originate within a specific window of B-cell differentiation, from lymphoid-primed multipotent progenitors (LMPPs), through common lymphoid progenitors (CLPs), to early B-cell progenitors [5]. This process is driven by the disruption of highly coordinated transcriptional programmes—governed by master regulators such as *IKZF1*, *EBF1*, and *PAX5* which normally ensure the orderly maturation of the B-cell lineage and the maintenance of B-cell identity [6]. The genomic landscape is defined by recurrent genetic alterations, where primary events establish the leukaemic subtype and secondary events show a non-random distribution across these genetic subgroups [7, 8]. This suggests genetic interactions, defined by the underlying molecular circuitry, where the cell states created by the initial first hit dictates the selective advantage provided by subsequent alterations [7].

Despite comprehensive cataloguing of the mutations driving BCP-ALL, and our understanding of how many of these mutated genes function in the normal B-cell maturation programme, a critical knowledge gap remains regarding how these elements integrate into a functional rubric. Therefore, we elected to develop an *in silico* model of B-cell maturation that would allow us to test the impact of introducing recurrent mutations in different combinations and at alternative developmental timepoints. Such a computational framework would allow us to decipher the underlying rules of leukaemogenesis and explain the molecular and phenotypic heterogeneity among patients. BCP-ALL is an ideal candidate for such modelling - while it exhibits inter-and intra-patient genetic heterogeneity, its overall genetic complexity is significantly lower than that of many solid tumours, being associated with a relatively small number of well-defined recurrent genetic abnormalities.

Most existing models of B-cell differentiation rely on qualitative methods, primarily Gene Regulatory Networks (GRNs) and Boolean frameworks [9–11]. However, none of these models studied BCP-ALL development nor fully accounted for the developmental window where BCP-ALL arises [5]. Furthermore, the binary nature of standard Boolean approaches restricts the ability to investigate copy number alterations, which are common in BCP-ALL. For these reasons, we selected a multi-valued logical formalism to develop our model. This approach uniquely balances computational tractability with the capacity to explicitly represent the non-binary, graded activity levels critical to capturing complex BCP-ALL genotypes. Mutations do not necessarily occur deterministically within the same cell stage of differentiation. Consequently, if modelled in such a way it may not be possible to reconstitute an appropriate evolutionary trajectory. Hence, a key feature of our model is the ability to introduce different genetic alterations stochastically i.e. at varying time points within deterministic simulations; allowing alternative paths to leukaemic transformation to be explored.

In this study, we first establish a model that faithfully recapitulates normal B-cell differentiation and subsequently use this to simulate the effects of archetypal genetic alterations driving BCP-ALL. Using the multi-valued logical framework implemented in the BioModelAnalyzer (BMA) [12], we describe the complex B-cell differentiation trajectory from the CLP to the Naive B-cell stage. We use this model to explore the genetic interactions driving leukaemogenesis, integrating the initiating mutation - fusion oncogene *BCR-ABL*, and simulating recurrent secondary events. We demonstrate that the clinical heterogeneity observed with respect to the stage of maturation arrest and aggressiveness of disease states can be explained by the combination, order and timing of secondary alterations. This model provides a platform to explore the genetic interactions underpinning BCP-ALL including clinically silent pre-cancerous states.

## 2 Methods

### 2.1 Logical Model Framework

The B-cell differentiation gene regulatory network (GRN) is formally represented as a directed graph *G* = (*V, E*). The set of nodes *V* = {*X*_1_, *X*_2_, … , *X_N_* } corresponds to the *N* key molecular components, including transcription factors, signalling molecules, and phenotypic markers. The set of directed edges E represents the regulatory interactions, categorized as either activating or inhibitory.

### 2.2 Model Building and Multi-valued Logic

We constructed a multi-valued logical model of the B-cell differentiation GRN. In this framework, each component *X_i_* is represented by an integer state variable *S*(*X_i_*) from a discrete set:

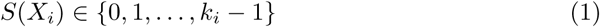

where *k_i_* is the number of distinct activity levels defined for node *X_i_*. A critical implementation detail in our model is the distinction between **constitutive genetic inputs** and **dynamic regulatory nodes**:

- **Gene Nodes:** For nodes representing genetic loci, *k_i_* = 3 were utilized to denote the static genotype status: inactive or homozygous deletion (0), heterozygous status (1), and normal/wild-type locus (2). These values function as constant parameters that constrain the regulatory logic and maximum activity levels of downstream functional nodes.
- **Dynamic Nodes:** Transcription factors, signalling components, and phenotypic markers (e.g., Apoptosis, Proliferation) evolve dynamically based on integrated regulatory signals.

Updates of the system are determined by a set of transition functions

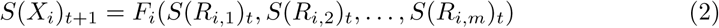

Models update synchronously; the values of all nodes are updated at each time step. The network was constructed and analysed within the BioModelAnalyzer (BMA). Stability analysis was used to assess prove the existence of a global fix point attactor.

An initial model was adapted from Collombet *et al.* [9]. The scope was refined to represent differentation states from the CLP through to the Naive B-cell stage. The model was expanded through a bottom-up approach, with interactions curated from 66 peer-reviewed publications (Supplementary Table 1).

### 2.3 State Classification and Scoring Functions

To map the high-dimensional state space of the simulation to discrete developmental stages, we implemented five multi-valued scoring functions. These functions act as automated classifiers:

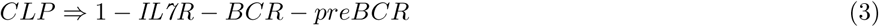

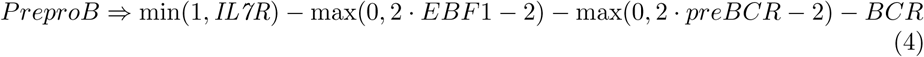

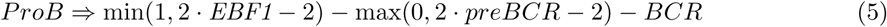

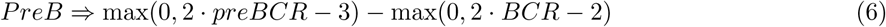

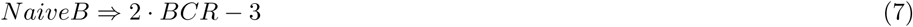

These functions integrate the activity levels of stage-specific markers to determine the cellular identity at each discrete time step *t*. Update functions were specified to ensure the stage definitions are mutually exclusive throughout the differentiation trajectory. The transition from CLP to Pre-pro-B (Eq.4) was characterized by the upregulation of IL7R activity [13]. Our logic correctly identified the Pro-B stage through the integration of *EBF1* signalling (Eq. 5) and the emergence of the pre-BCR complex [14]. We further confirmed the discriminatory power of the Pre-B and Naive B scoring functions; the transition to the Pre-B stage was accurately captured by the peak in pre-BCR scoring (Eq. 6), while the Naive B state was strictly defined by complete BCR expression (Eq. 7) [15].

### 2.4 Fitness score

To quantify the pathological aggressiveness of each simulated genetic profile, we developed a composite phenotypic fitness score (Φ). This metric integrates four key cellular parameters—**Proliferation** (*P*), **DNA Damage** (*D*), **Apoptosis** (*A*), and **Cell-Cycle Arrest** (*C*)—into a single quantitative value designed to reflect the oncogenic potential and survival advantage of the leukaemic clones. Given the multi-valued logic of our framework, these phenotypic markers operate within a discrete range of [0, 4]. The fitness score is defined as follows:

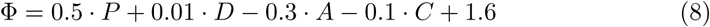

The coefficients were assigned to weight the relative contribution of each metric to the leukaemic phenotype, prioritizing proliferative capacity and survival. To ensure a non-negative output (Φ ≥ 0) across the entire state space, a constant of 1.6 was included; this value specifically compensates for the maximum possible negative contribution from the inhibitory terms (−0.3 · 4 − 0.1 · 4 = −1.6).

### 2.5 Model Validation

Our model represents a fixed and sequential developmental process. To verify its correctness, we proved the stability of the model in different environments at the start or end of differentiation. A proof of stability here proves that there exists a single global, fixpoint attractor and no cycles, representing a unique developmental end-state. We assessed the stability of the system at the initial state (CLP) in the absence of exogenous inputs (*IL7* and *IKZF1*) (Supp Fig.1)) and at the terminal differentiation state (Supp Fig.2). The observed stability of these fixed points, combined with the strictly sequential order of TF expression during differentiation, confirms that a single trajectory accurately represents the biological behaviour.

The physiological relevance of the model was established through validation against biological literature under two conditions. First, developmental states identified in the unperturbed network (*BCR*-*ABL* = 0) were compared to characteristic molecular profiles of healthy B-cell precursor stages. Second, the model’s dynamic responses were tested via *in silico* perturbations.

Previous work has interpreted individual patient leukaemia datasets in terms of likely stages of developmental arrest [7, 16]. To test our model’s ability to reproduce these observations, the perturbed model (*BCR*-*ABL* = 1) had a second hit introduced with the KO of *IKZF1, EBF1* and *PAX5*. These alterations, represented in the model by specifying an update function equal to either 1 or 0 depending on the specific gene, were specified at the start of the simulation and the stage of arrest recorded and compared to patient profiles.

### 2.6 Simulations

All computational simulations were performed using the pyBMA library^1^, a Python-based interface for the BMA engine. To explore the evolutionary landscape of *BCR-ABL* positive BCP-ALL, we executed a systematic series of *in silico* perturbations. We defined a set of 784 unique evolutionary trajectories, representing all possible permutations of the four most common secondary alterations: *IKZF1* (homozygous and heterozygous), *EBF1* (heterozygous), *PAX5* (heterozygous), and *CDKN2A* (homozygous). For each simulation, the *BCR-ABL* oncogene node was initialized at a value of 0 for WT and 1 for oncogenic to represent an initiating event. Secondary hits were introduced at specific developmental windows (CLP, Pre-pro-B, Pro-B, or Pre-B) by overriding the target node’s logical function with a constant value corresponding to the alteration type (e.g., setting a node to 0 for a homozygous deletion).

Each simulation was run for a minimum of 30 steps—a duration found to be sufficient for the WT model to reach a stable B-cell attractor—and extended when limit cycles (cyclic attractors) exhibited long periodicity. We recorded the full state-space history of master regulators and phenotypic markers at every time step. Trajectories were subsequently filtered for developmental reachability; a trajectory was deemed unreachable if a secondary alteration was programmed to occur at a developmental stage that the cell could no longer attain due to a prior maturation arrest (e.g., an alteration programmed for the Pre-B stage following a Pro-B arrest). To identify the 58 highly probable trajectories, we applied a stepwise optimization filter. An alteration sequence was conserved only if each successive genetic hit resulted in a higher phenotypic fitness score (Φ) compared to the previous state. This represents selection of clones with superior survival or proliferative advantages.

### 2.7 Data Availability

The source code for the multi-valued logical model of B-cell differentiation and the scripts used for the systematic *in silico* perturbation analysis are hosted on Github (https://github.com/MBougueon/ModALL) and Zenodo (https://doi.org/10.5281/zenodo.20135289). A web-based interactive dashboard of the model is available at http://bcp-all.biomodelanalyzer.org/, allowing users to replicate the simulations and explore the impact of specific genetic perturbations on B-cell maturation.

## 3 Results

### 3.1 Regulation of B-cell differentiation by a core pentad of transcription factors

To investigate the relationship between shared processes governing leukaemic evolution, we first developed a qualitative model of healthy B-cell differentiation. The interaction network and update rules were specified by curating the literature describing the regulatory relationships governing B-cell differentiation and their perturbation in BCP-ALL, using computational mining supplemented by large language models (LLMs), and incorporating interactions of five key transcription factors *IKZF1*, *EBF1*, *E2A*, *PAX5*, and *FOXO1* from published models [9, 17]. This framework was specifically constrained to span the developmental window where BCP-ALL-associated developmental arrest occurs: from the CLP stage through the Pre-pro-B, Pro-B, and Pre-B stages, terminating at the Naive B-cell stage [5, 16] Fig.1. To ensure high model fidelity, identified interactions were stringently filtered based on experimental evidence from relevant cell systems and cross-referenced against curated databases such as KEGG [18] (Supplementary Table 1).

**Fig. 1:**
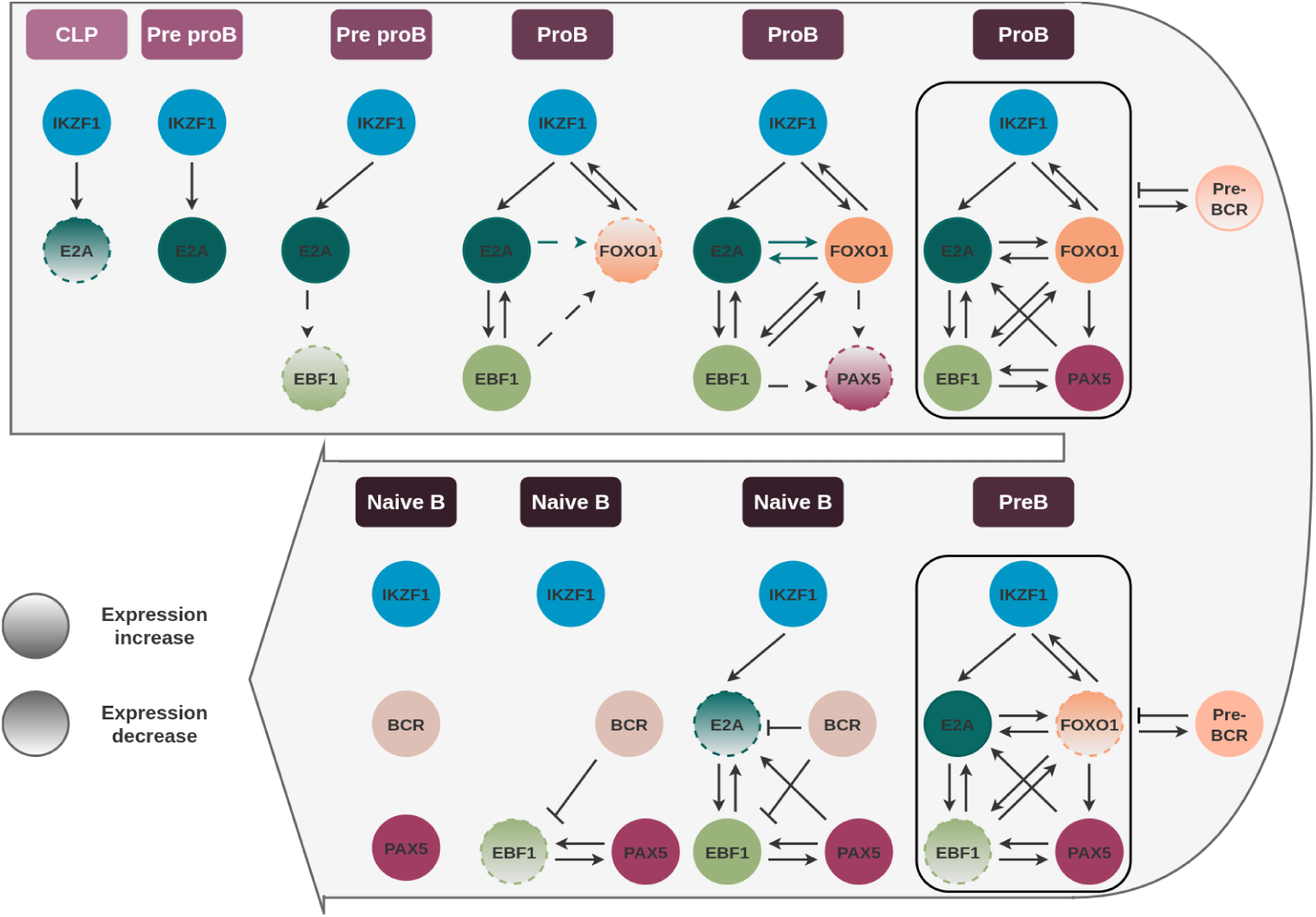
Regulatory Dynamics and Phenotypic Transitions in Healthy B-cell Differentiation. Schematic representation of the core regulatory network involving five key transcription factors (TFs): *IKZF1*, *E2A*, *EBF1*, *FOXO1*, and *PAX5*. The diagram illustrates the hierarchical induction and feedback loops governing the transition from CLP to naive B-cells.

The initiation of the B-cell program is spearheaded by *IKZF1*, which is expressed prior to the CLP stage [19]. *IKZF1* facilitates the induction of *E2A*, primarily through the repression of the myeloid-priming factor *CEBPA* [20]; notably, while this repression is a critical biological priming step, it is simplified to *IKZF1* inducing *E2A* directly in our current model. Following the expression of *E2A*, the expression of the Interleukin-7 receptor (*IL7R*) is induced [21]. Signalling through the *JAK-STAT* pathway subsequently triggers the activation of *EBF1* [22], which serves as a central scaffold for the B-lineage by promoting the expression of *FOXO1* and *PAX5* [21, 23–26]. Ultimately, the synergistic activity of these five master regulators drives the assembly and expression of the pre-B-cell receptor (pre-BCR) and the mature BCR—a critical milestone marking the commitment to B-cell identity [17]. Notably, this developmental transition, and the regulatory checkpoint it represents, was absent from previous modelling efforts.

The inherent complexity of several signalling pathways implicated in B-cell development—specifically *JAK-STAT*, *MAPK*, *PI3K*, and *WNT* —necessitated their abstraction into discrete representative nodes. By representing these cascades as simplified logical nodes, we maintain model tractability while preserving the essential signalling inputs required for lineage progression. As mutations in BCP-ALL are typically heterozygous or homozygous deletions, we built our model to be sensitive to gene dosage, implementing a multivalued logic using activity levels from 0 (homozygous loss) to 2 (wild-type) for each regulatory node (see Materials & Methods 2.2). The key phenotypic parameters of Proliferation, Apoptosis, Cell-Cycle Arrest, and DNA Damage (Fig.2 C) are implemented as output nodes regulated by well-characterized gene families such as *CDKN*, *CCND*, and *BCL*. These phenotype nodes can have values between [0 : 4] allowing a more fine-grained tracking of phenotypic evolution during differentiation. All the interactions encoded in the model are visualized in Sup Fig 3.

**Fig. 2:**
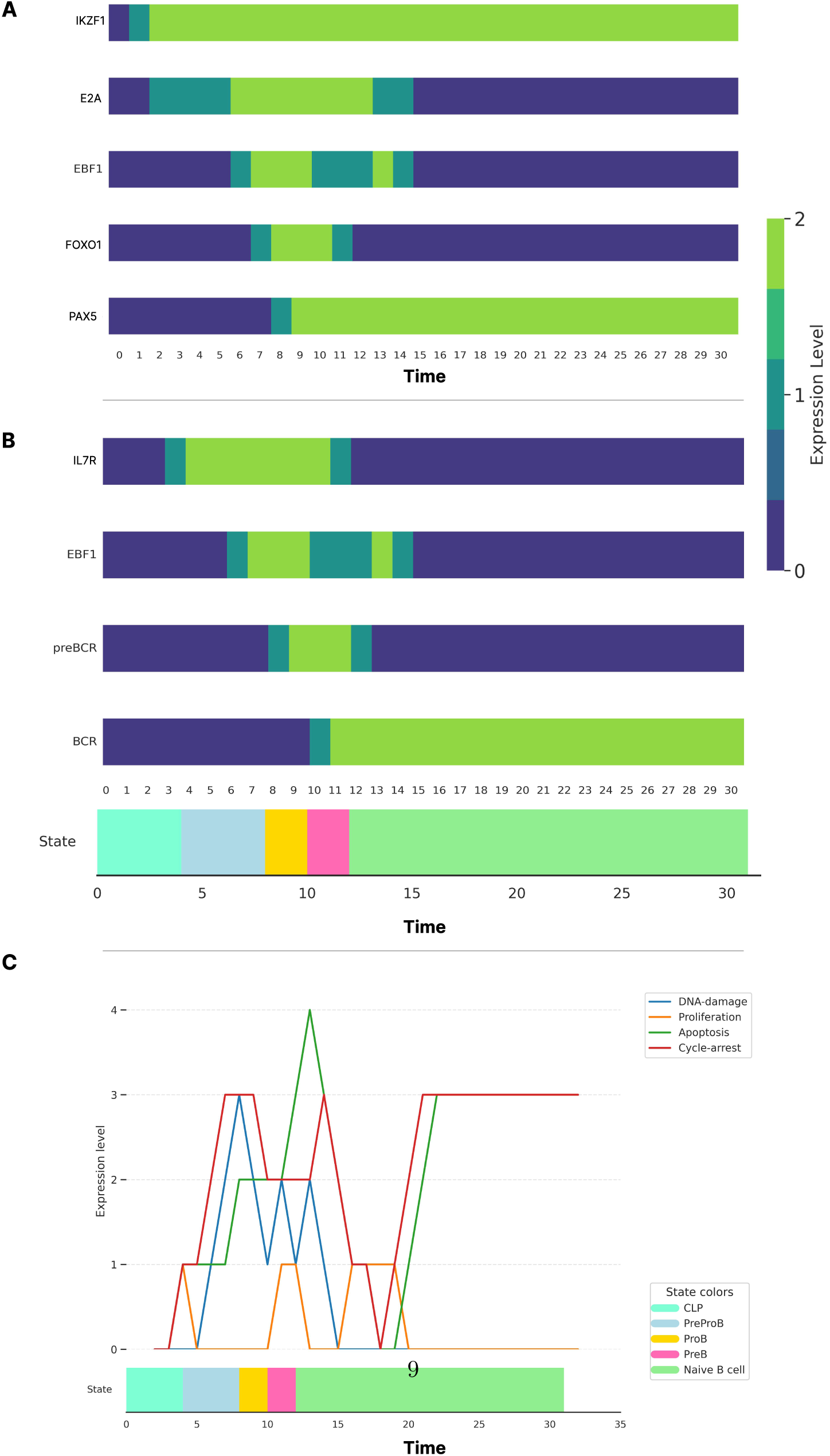
Regulatory Dynamics and Phenotypic Transitions in Healthy B-cell Differentiation. **A**. Heatmap of simulated transcription factor activity. The temporal profiles recapitulate the sequential activation observed *in vivo*, notably capturing the transient down-regulation and subsequent recovery of *EBF1* levels during the pro-B to pre-B transition. **B**. Heatmap of stage-specific marker activity. The simulation demonstrates the dynamics of nodes used to define distinct cellular states, highlighting the sequential progression through the B-lineage program. **C**. Temporal simulation curves of phenotypic markers. The top panel displays the predicted levels of Apoptosis, Proliferation, Cell-Cycle Arrest, and DNA Damage. The bottom panel provides a differentiation timeline, demarcating the transitions between the CLP, Pre-pro-B, Pro-B, Pre-B, and Naive B stages as defined by the scoring functions.

To model cell state transitions we considered both the initial CLP state and final naive B-cell state as stable global fixed point attractors. Simulations were performed by setting all node values to 0 and initiating *IKZF1* expression and exposure to *IL7* (*IL7* and *IKZF1* nodes = 2) which activate *E2A* and *IL7R*, triggering the differentiation process. Node values were tracked over 5 discrete stages and their activation sequence compared to the known sequence of events in B-cell differentiation. Our model reproduces the sequential activation of the core transcription factors (Fig.2 A) although this behaviour was not explicitly programmed.

Furthermore, our model produced complex dynamics of *EBF1* expression, consistent with experimental evidence ([27], Fig. 2A-B) and providing novel mechanistic insight. The transient downregulation post-peak is driven by pre-BCR signalling, which temporarily antagonizes *EBF1* activity. This is subsequently compensated for by *BACH2*, a transcription factor induced by *PAX5* that stabilizes the B-cell program [14], ensuring lineage commitment.

Finally, we compared phenotypes captured by the model to known behaviours to ensure its biological soundness. B-cell differentiation is intrinsically coupled to the regulation of proliferation and survival, centering on the formation of a functional pre-BCR during the transition from Pro-B to proliferating Pre-B cells. *EBF1*, *PAX5*, and *IKZF1* induce the formation of pre-BCR through *RAG1*, *VPREB1* and *IGLL1* expression. In turn, this induces *MYC* -mediated proliferation [14, 28], and release of cell-cycle arrest imposed by *BCL6* ’s repression of the *CDKN* family [29, 30]. We used the combinatorial pattern of *IL7R*, *EBF1*, *pre-BCR* and *BCR* to distinguish the differentiation stages in our model (Fig 2B). This captured the expected sequence of events during the transition from CLP, through pre-pro-, pro- and pre-B to naive B-cell. At the pre-B stage, increased proliferation and decreased cell cycle arrest are observed as expected (Fig 2C). Continued pre-BCR signalling ultimately promotes progression to non-proliferative Small Pre-B cells (early Naive B in the model, [27]). Our model also captures this with pre-BCR expression reducing *EBF1* activity and *JAK-STAT* signalling, leading to a reduction in *CCND3* expression and a decrease in proliferation (Fig 2B, C).

### 3.2 Model recapitulates anticipated transcription factor dependencies

Our model recapitulates the process of B-cell differentiation through coordinated action of the core transcription factors (Fig 1). Experimental evidence from cell lines and mouse models shows that loss-of-function mutations of the core transcription factors leads to differentiation arrest at different points in B-cell development [31–34]. We used this information to test the performance of our model, simulating TF knockouts by setting the value of the respective nodes to 0. Knockout of any single TF disrupted the sequential activation of the network (Fig. 3A), leading to the loss of downstream TF activity and ultimately failure to assemble the mature B cell receptor and reach the naive B cell stage (Fig. 3 B & C) [15, 32]. Consistent with the literature [19, 35–37], *IKZF1* KO resulted in failure to initiate *E2A* expression and in turn activate the downstream TFs while knockout of *E2A* itself resulted in failure of downstream activation with *IKZF1* unaffected (Fig. 3A). The model also recapitulates the specific dependency of *IL7R* on *E2A* and its upstream regulator *IKZF1* Fig. 3B [20, 21], with loss of either gene resulting in differentiation arrest at the CLP stage Fig. 3C, in agreement with experimental observations [31, 38]. Note that in experimental models, *IKZF1* knockout results in arrest at the earlier multipotent progenitor (MPP) stage but the earliest state described in our model is CLP. *EBF1* knockout in our model results in failure to fully activate *FOXO1*, no *PAX5* activation, and ultimately differentiation arrest at the PreProB stage with failure to express the pre-BCR, consistent with experimental observations [19, 32] (Fig. 3A-C). *PAX5* knockout results in differentiation arrest at the preB stage consistent with its described role as a gatekeeper of pre-B to naive B transition [34]. A novel prediction of the model is that the *EBF1* knockout results in failure to down-regulate the *IL7R* due to the absence of preBCR-mediated silencing of *IL7R* directly and through the transient antagonism of *EBF1* Fig. 3B.

**Fig. 3:**
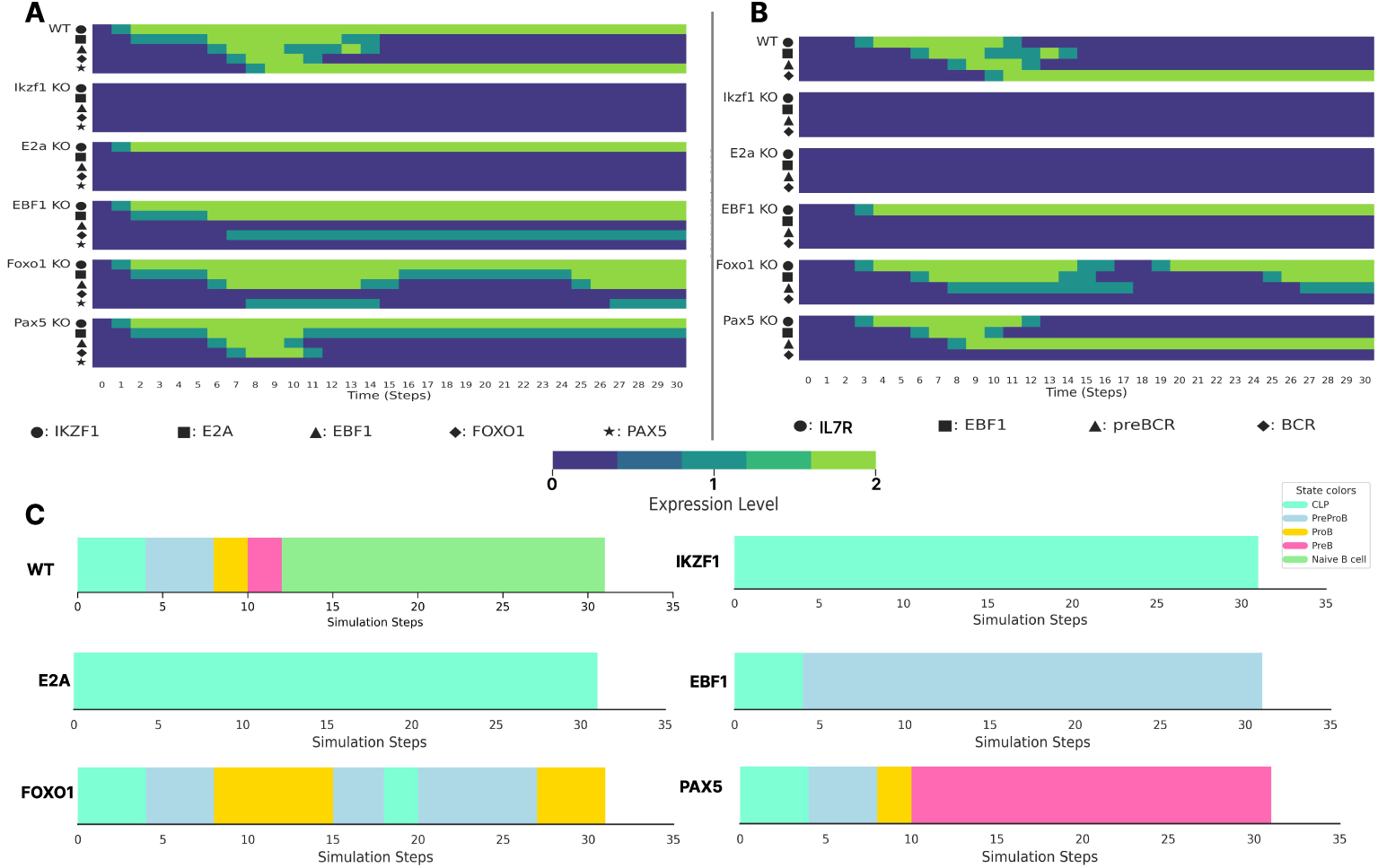
Sensitivity analysis of the model following individual TF alterations. Each of the five core transcription factors (IKZF1, *E2A*, *EBF1*, *FOXO1*, and *PAX5*) was inactivated to assess network robustness and differentiation dependency. **A**. Heatmap of simulated transcription factor activity profiles. Loss of individual transcription factors delays or prevents activation of other genes, visible as a right shift in bright colours or loss of activity (dark blue). **B**. Heatmap of stage-specific marker dynamics under KO conditions. The simulation illustrates how individual TF removals perturb the expression of nodes used to define cellular states, demonstrating the differential requirement of each factor for stage-specific marker induction. **C**. Simulated differentiation time-line. The timeline demarcates the impact of each TF knockout on the transitions between the CLP, Pre-pro-B, Pro-B, Pre-B, and Naive B stages. These results highlight the specific developmental checkpoints where maturation arrest is induced by the absence of each regulator.

Intriguingly, the knockout of *FOXO1* resulted in a cyclic attractor (SFig. 4). This oscillation drives the cell back and forth between the CLP and Pro-B differentiation stages. This suggests that while the initial signals for B-cell commitment are present, *FOXO1* is the essential molecular switch required to commit the cell to a stable Pro-B identity. Without it, the system fails to sustain the B-lineage program and perpetually cycles back to a more primitive progenitor state. This “stutter” accurately reflects experimental observations that *FOXO1* -deficient progenitors fail to effectively transit the Pro-B checkpoint (Fig. 3C) [39].

The behaviour of our model demonstrates a high degree of concordance with experimental results and accurately recapitulates complex wild-type dynamics, including non-trivial phenomena such as *EBF1* transient oscillations, while the differentiation endpoints observed in the TF knockout simulations agree with established *in vitro* and *in vivo* data [15, 32].

### 3.3 *BCR-ABL* accelerates but does not arrest differentiation

The t(9;22) translocation forms the *BCR-AB* L fusion gene, is the initiating event in approximately 4% of paediatric and 25% of adult BCP-ALL cases, and is associated with poor clinical outcomes. To explore its impact on our model, *BCR-ABL* was integrated into the network as a pleiotropic node that modulates the JAK-STAT pathway and *BCL6*, while indirectly inhibiting *BACH2* via the repression of *PAX5* [29, 40–42] (Fig. 4A). In our model, *BCR-ABL* activity alone did not trigger the B-cell differentiation program. However, once the differentiation program is initiated, *BCR-ABL* accelerates lineage progression by reducing the duration of the Pre-Pro-B phase and reducing the number of simulation steps to complete differentiation (Fig. 4B). This acceleration is driven by the constitutive activation of *JAK-STAT* signalling, which facilitates the premature induction of *EBF1* and other transcription factors (Fig. 4C). Furthermore, BCR-ABL activity results in sustained expression of E2A which may affect B-cell maturation.

**Fig. 4:**
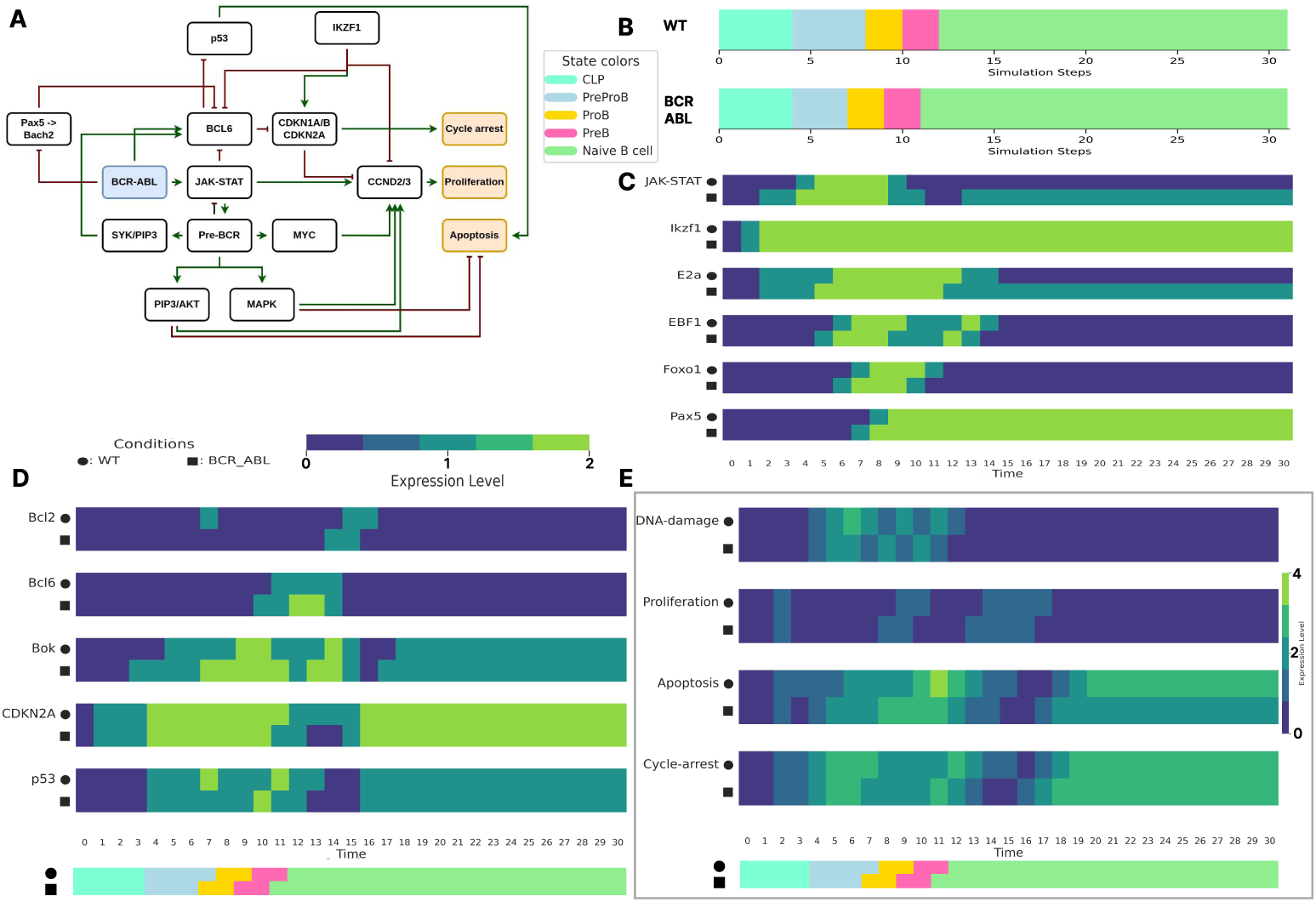
Impact of *BCR-ABL* signalling on B-cell regulatory dynamics and differentiation progression. **A**. Schematic representation of the integrated regulatory network. The diagram illustrates the mechanistic influence of *BCR-ABL* on key phenotypic nodes, specifically detailing its role in the evasion of Apoptosis, the induction of Proliferation, and the bypass of Cell-Cycle Arrest checkpoints. **B**. Simulated differentiation timeline. The timeline demarcates the impact of *BCR-ABL* on the transitions between the CLP, Pre-pro-B, Pro-B, Pre-B, and Naive B stages. These results highlight an acceleration of the differentiation process under *BCR-ABL* alteration, suggesting a loss of temporal control during early B-cell commitment. **C**. Comparative heatmap of transcription factor expression dynamics. A side-by-side simulation of WT versus *BCR-ABL* conditions. **D**. Comparative heatmap of the most altered gene expression dynamics. A side-by-side simulation of WT versus *BCR-ABL* conditions. **E**. Comparative heatmap of phenotypes dynamics. A side-by-side simulation of WT versus *BCR-ABL* conditions. The heatmap highlights the impact of *BCR-ABL* on the phenotypic profile of a preleukaemic B-cell.

*BCR-ABL* is also expected to have a systemic effect on the network and cellular phenotypes. Our model shows that several pro- and anti-apoptotic genes, and cell cycle regulators, exhibit altered activity in response to *BCR-ABL* (Fig. 4D). The shift in dynamics broadly correlates with the accelerated PreProB phase, while increased *BOK* and *BCL6* levels have opposing impacts on *P53* -mediated apoptosis, and reduced *EBF1* leads to a reduction in cycle arrest as the simulation transitions through proB and preB stages (Fig. 4D, E). In the latter stages of differentiation, *BCR-ABL* maintains JAK-STAT activity, escaping the physiological down-regulation typically mediated by the pre-BCR and BCR pathways [14, 29], reducing apoptosis levels (Fig. 4C, E). During the Pre-B stage, we observed a down regulation of *P53*, driven by the BCR-ABL-mediated inhibition of *BACH2*, a known repressor of the *P53* locus [30]. This suppression of the *P53* -dependent DNA damage response, combined with the up regulation of *BOK* (Fig. 4D), creates a permissive environment for the survival of progenitors that would otherwise be eliminated by stringent developmental checkpoints.

This genetic alteration leads to an overall increase in clonal fitness, offering and explain for the positive selection of *BCR-ABL*. This arises primarily from the simultaneous promotion of survival signalling and the silencing of tumor-suppressive checkpoints. We observe a sustained decrease in Apoptosis levels (Table 1), and Proliferation rises occur earlier in development while maintaining similar overall levels (Fig. 4E). These simulations predict that *BCR-ABL* acts as a catalytic driver that compresses the early windows of B-cell commitment via accelerated *JAK-STAT* and *EBF1* kinetics. They further suggest that BCR-ABL alone does not induce differentiation arrest, consistent with clinical observations in which patients invariably have additional mutations, often in core components of the B-cell differentiation machinery [7, 16].

**Table 1:** WT vs BCR-ABB, phenotypic marker mean per transition states and weighted average across all simulation steps.

|  | Proliferation | Apoptosis | Cycle-Arrest |
| --- | --- | --- | --- |
| CLP | 0.33/0.33 | 0.667/0.33 | 0.667/0.667 |
| Pre-ProB | 0/0 | 1.5/1.667 | 2.75/2.75 |
| ProB | 0.5/0.5 | 2/2.5 | 2/2 |
| PreB | 0.5/0.5 | 3.5/3 | 2/2 |
| Naive B Cell | 0.21/0.2 | 2.26/1.7 | 2.42/2.4 |
| Average | 0.30876/0.30666 | 1.98598/1.84 | 1.96756/1.94668 |

### 3.4 Secondary Hits Drive Stage-Specific Differentiation Arrest

The preleukaemic state caused by *BCR-ABL* and characterized by rapid lineage progression and impaired surveillance might provide a fertile ground for the acquisition of secondary genetic lesions, ultimately leading to maturation arrest and the clonal expansion that is the hallmark of clinical BCP-ALL. In *BCR-ABL* patients, loss of *IKZF1*, *PAX5*, *EBF1*, and *CDKN2A* are frequent secondary mutations [7, 16]. It has been proposed that the stage of developmental arrest is determined by the specific loss of different core transcription factors. *EBF1*, *IKZF1*, and *PAX5* losses are associated with arrest at the Pre-pro-B, Pro-B, and Pre-B stages, respectively [16, 43] (Fig. 5A). To explore how mutations combine, we created secondary alterations in a *BCR-ABL* background and compared the model with clinical observations.

**Fig. 5:**
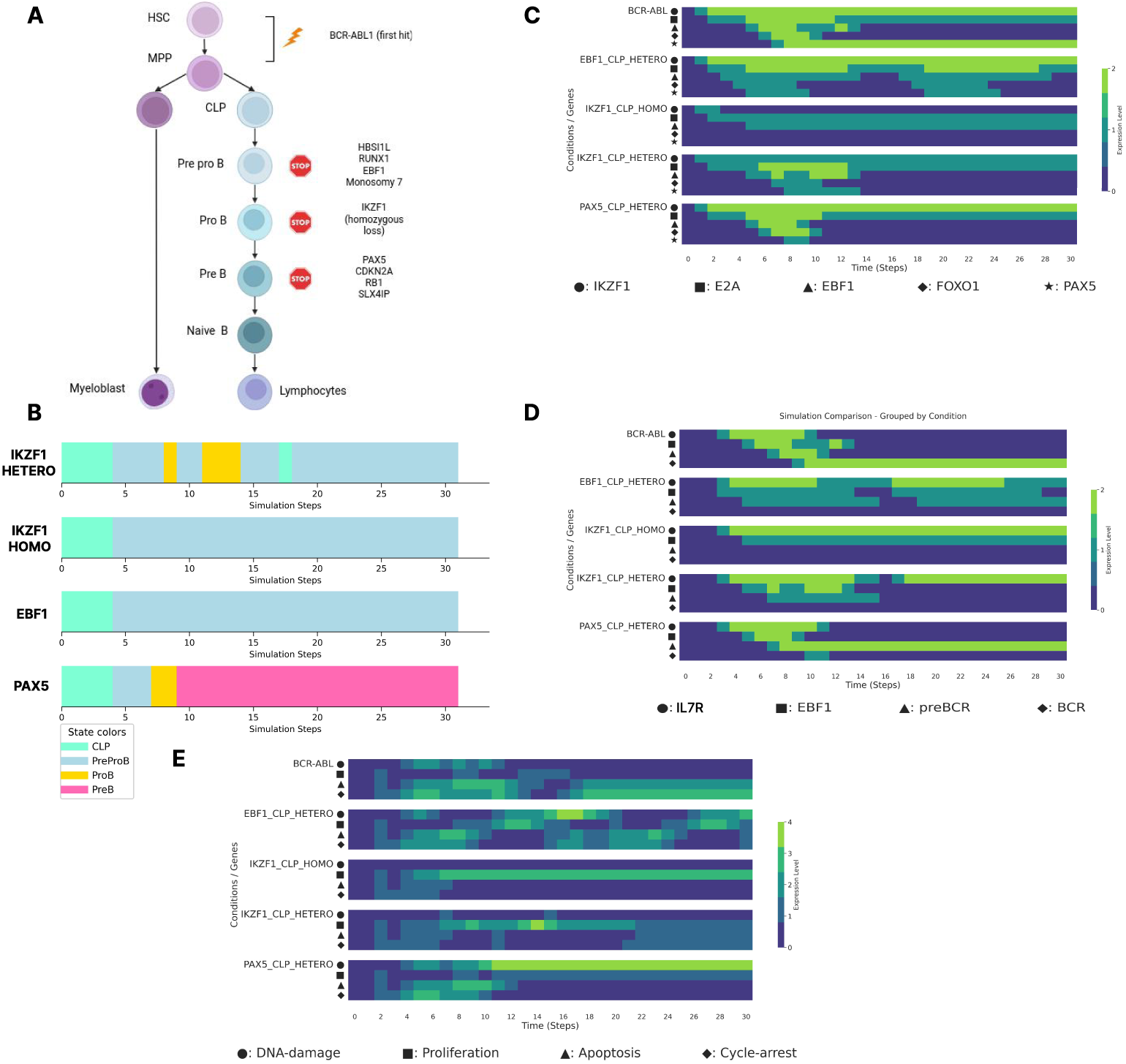
Cooperative effects of *BCR-ABL* and patient-specific deletions on B-cell differentiation arrest. **A**. Schematic of *BCR-ABL* and secondary hits in BCP-ALL. The diagram illustrates the integrated influence of *BCR-ABL* signalling and secondary hits (e.g., *IKZF1*, *EBF1*, or *PAX5* deletions) on B-cell commitment. The schematic highlights how specific secondary lesions dictate the precise developmental stage at which differentiation arrest occurs. Adapted from Kim *et al.* [16] and created with BioRender. **B**. Simulated differentiation kinetics and maturation arrest checkpoints. The timeline demarcates the impact of *BCR-ABL* in combination with secondary hits, including heterozygous *EBF1* deletion, homozygous and heterozygous *IKZF1* deletion, and heterozygous *PAX5* deletion. The results demonstrate distinct stage-specific blocks: arrest between the Pre-pro-B stages for *EBF1* loss, at the Pre-pro-B stage for *IKZF1* loss, and at the Pre-B stage for *PAX5* loss. These computational predictions are concordant with clinical observations by Kim *et al.* [16] and Iacobucci *et al.* [43]. **C**. Comparative heatmap of transcription factor expression dynamics. A side-by-side simulation of *BCR-ABL* alone and with heterozygous *EBF1* deletion, homozygous or heterozygous *IKZF1* deletion, or heterozygous *PAX5* deletion conditions. **D**. Comparative heatmap of stage-specific marker dynamics. Evolution of lineage-defining markers across the same genetic conditions described in (C). **E**. Comparative heatmap of phenotypic d^15^ynamics. Temporal evolution of Proliferation, Apoptosis, Cell-Cycle Arrest, and DNA Damage scores under the same genetic conditions described in (C).

We initially simulated homozygous (-/-) and heterozygous (+/-) *IKZF1*, heterozygous *EBF1*, and heterozygous *PAX5* deletion at the CLP stage in our *BCR-ABL* model. This resulted in maturation arrest at the pre-pro-B stage for *IKZF1* (-/- and +/-) and *EBF1* (+/-) and at the pre-B stage for *PAX5* (+/-) Fig.5 B, consistent with observations in patients 5 A [16]. Interestingly, *IKZF1* (-/-) loss in the context of *BCR-ABL* results in later arrest than in the wild-type context, occurring at the pre-pro-B stage, as opposed to the CLP stage. This results from *BCR-ABL* induction of *E2A*/*EBF1* through *JAK-STAT*, effectively reducing the dependency on *IKZF1*.

While several genetic perturbations implemented at CLP result in arrest at the same developmental stage, the terminal state alone does not capture the underlying molecular heterogeneity between leukaemic profiles. Our model reveals that heterozygous *EBF1* loss results in the emergence of a cyclic attractor spanning the Pre-pro-B stage (Fig. 5C). This oscillation arises from the activity of downstream nodes, notably *PAX5* and *FOXO1*, and results in a cyclic phenotypic profile (Fig. 5E), where periods of high Proliferation and DNA Damage accumulation are followed by transient intervals of Cell-Cycle Arrest and Apoptosis

Similarly, the dosage of *IKZF1* produces distinct molecular profiles: the heterozygous deletion of *IKZF1* results in a more advanced Pre-pro-B state compared to homozygous, evidenced by the sustained expression of *EBF1* and a broader repertoire of activated transcription factors (Fig. 5C,D). These findings accord with clinical data describing alternative groups of BCR-ABL patient groups, with homozygous loss resulting in intermediate proB arrest while heterozygous loss collaborates with additional mutations to give early or late proB arrest [16]. Interestingly, the homozygous *IKZF1* deletion has a more aggressive phenotypic profile characterized by a complete absence of Apoptosis and Cell-Cycle Arrest, coupled with sustained high levels of Proliferation (Fig. 5E). The *PAX5* heterozygous deletion similarly displays an absence of Apoptosis and Cell-Cycle Arrest, yet it is distinguished by a lower baseline of Proliferation alongside potentially high levels of DNA Damage (Fig. 5E). This suggests a state of genomic instability that lacks the rapid clonal expansion characteristic of *IKZF1* mutants, potentially explaining the different leukaemogenesis trajectories associated with these secondary hits.

The sensitivity to mutations occurring at or prior to CLP raises the question of whether this impact is dependent on the stage of differentiation. To explore this relationship we performed simulations with 17 clinically observed combinations of the 4 secondary alterations (Fig. 6 A) [16], in all possible sequences, and compared them with respect to the resulting state of maturation arrest. We find that the timing and order of alterations are as critical as the combination of alterations. A single genotype can lead to divergent outcomes, including stage of differentiation arrest, depending on the developmental windows in which the different mutations occur (Fig. 6B, top panel). Distinct genetic profiles can also converge onto identical differentiation arrest points, provided they occur within specific developmental windows (Fig. 6B, bottom panel).

**Fig. 6:**
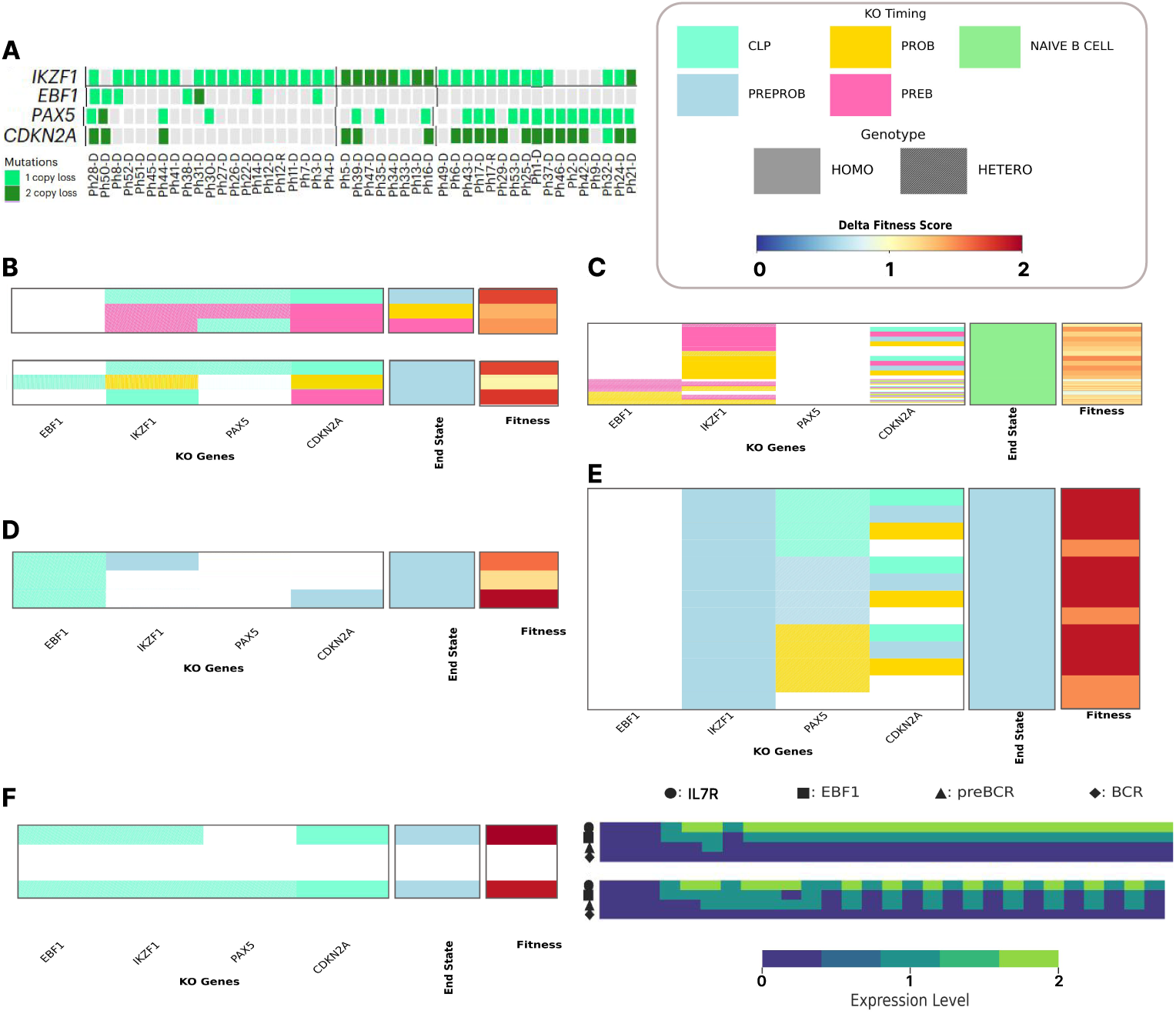
Characterization of secondary hit hierarchies and evolutionary trajectories in *BCR-ABL* BCP-ALL. **A** Genetic profile of *EBF1, IKZF1, PAX5* and *CDKN2A* in *BCR-ABL* patient. Adapted from [16]. **B–E**. Comparative landscape of genetic alterations and phenotypic outcomes in *BCR-ABL* BCP-ALL. Each heatmap summarizes simulated clinical profiles across four dimensions: **Initial Hit Hierarchy**: The primary regulatory perturbation driving the lineage. **Combinatorial Alteration Burden**: The first column indicates the most frequent clinical combinations involving *EBF1*, *IKZF1*, *PAX5*, and *CDKN2A* alterations, reflecting the complexity of cumulative genetic lesions. **Maturation Arrest Point**: The second column displays the predicted final differentiation state at the simulation terminus, identifying the specific developmental checkpoint (e.g., Pro-B, Pre-B) where the lineage is blocked. **Oncogenic Fitness Score**: The final column provides a composite phenotypic fitness score, calculated as the weighted mean of Apoptosis, Proliferation, Cell-Cycle Arrest, and DNA Damage profiles. This score serves as a quantitative proxy for the clinical aggressiveness of each specific genetic trajectory. **B**. Divergence and convergence of genetic alterations **C**. Combinations reaching the Naive B state, representing incomplete maturation blocks. **D**. Early Arrest Landscape: Impact of *IKZF1* and *CDKN2A* alterations on the Pre-pro-B arrest phenotype in the context of *EBF1* disruption. **E**. Late Arrest Landscape: Evolutionary trajectories involving *PAX5* alterations and the sequential requirements for maximizing fitness in Pre-B arrest phenotypes. **F**. Comparative heatmap of stage-specific ma^1^r^8^ker dynamics. Evolution of lineage-defining markers across the different genetic conditions showing circular attractor state.

This has a range of implications for BCP-ALL carcinogenesis. Firstly, a subset of combinations can progress to the Naive-B state, avoiding a complete maturation block. This occurs when the loss of *EBF1* or *IKZF1* occurred after the Pre-pro-B stage in the absence of *PAX5* loss, or when *PAX5* was lost after the Pre-pro-B stage without early *EBF1/IKZF1* deletions (Fig.6 C). Since maturation arrest prior to the Naive-B stage is a hallmark of BCP-ALL, these trajectories are inconsistent with leukaemogenesis and were therefore excluded from further analysis. Secondly, all other combinations result in arrest at the Pre-pro-B or Pre-B stages, aligning with immunophenotypes commonly observed in BCP-ALL as described by Kim <u>et al.</u> [16] and Iacobucci <u>et al.</u> [43]. Early loss of *EBF1* or *IKZF1* consistently led to a Pre-pro-B arrest, a phenotype unaffected by *PAX5* status (Fig.6 E Supp Fig6). Mechanistically, our model explains that later differentiation stages can only be reached with normal *EBF1* expression, or late alteration, and retention of at least one functional copy of *IKZF1* until the late Pro-B stage. Without these conditions, the activation threshold for *PAX5* is never met, halting the program prematurely. Notably, while Pre-B arrest is frequently linked to *PAX5* deletions [34], our model confirms that a Pre-B block can occasionally occur without *PAX5* loss due to cumulative network disruption.

Finally, we sought to link our model’s phenotypes to clinical severity. To achieve this we propose an evolutionary fitness score (*ϕ*), calculated as a weighted mean of cell phenotypes directly relevant to selection: Apoptosis, Proliferation, Cell-Cycle Arrest, and DNA Damage, with Proliferation and DNA damage given a positive coefficient and the others negative. We measure *ϕ* either as the average across the trajectory or on the last 5 steps of the simulations (Supp Fig.7,6). When grouping trajectories by end-state, we observed that our fitness score is highly sensitive to alteration context. In the Pre-pro-B group, *IKZF1* loss was associated with higher fitness than *EBF1* loss (Fig.6 D, Supp Fig6), primarily through the downregulation of *CCND2/3* and *BCL6* and the reduction of *CDKN* family expression, increasing proliferation and reducing apoptosis regardless of stage of developmental arrest. These findings align with the well-documented clinical association between *IKZF1* deletions (often referred to as the Ikaros-plus phenotype) and poor prognosis, characterized by increased chemoresistance and lower overall survival rates in *BCR-ABL* BCP-ALL patients [7, 16, 43–46]. A similar observation was made for *CDKN2A*; our model shows that the earlier this loss occurs, the more impact on overall fitness (Fig.6 D, Supp Fig6).

Beyond allowing us to filter evolutionary trajectories based on expected clonal survival, the inclusion of fitness in our model allows us to visualise the diverse outcomes of different effects of mutations in terms of intersecting impacts of fitness (and phenotypes) and differentiation state (Fig.6 F, Supp. Fig. 8, 9, 10, 11). This reveals that the clinical heterogeneity can be explained by the combination, order and timing of secondary alterations. The presence of functional *EBF1* leads to homogenous molecular profiles, whilst mutation of the gene is associated with highly heterogeneous outcomes and is sensitive to the specific timing of mutation. For example, where *EBF1* loss and heterozygous *IKZF1* alterations occur at different developmental stages, the resulting circular attractors differ in their periodic behaviour and gene expression dynamics depending on the precise timing of the hits (Supp. Fig. 8, 9). This suggests that the order of alteration in the evolution of the clone doesn’t just change the fitness of the cell, but can fundamentally alter the stability of the leukaemic state, shifting it from a static arrest to a dynamic, oscillating regulatory cycle.

This visualization reveals that some trajectories that can be simulated are not possible in leukaemogenesis. For example, if *BCR-ABL* and *EBF1* alterations at the CLP stage induce a maturation arrest at the Pre-pro-B stage, subsequent alterations programmed for the Pre-B window become unreachable. This allows us to reduce 748 simulatable trajectories to 147 that are biologically feasible (Supp Fig 6). From the feasible subset, we identified 58 highly probable evolutionary trajectories by applying an evolutionary filter based on fitness (Supp Fig. 12). This dual-filtering approach—accounting for both developmental reachability and stepwise fitness optimization— allowed us to map the probable evolutionary paths driving the transition from healthy progenitors to malignant BCP-ALL populations (Fig. 7). These 58 evolutionary trajectories converge onto the 17 distinct genetic profiles observed clinically, suggesting a mechanistic explanation for the heterogeneity found among patients, even those presenting with similar genetic profiles at the point of diagnosis (Fig.6 A).

**Fig. 7:**
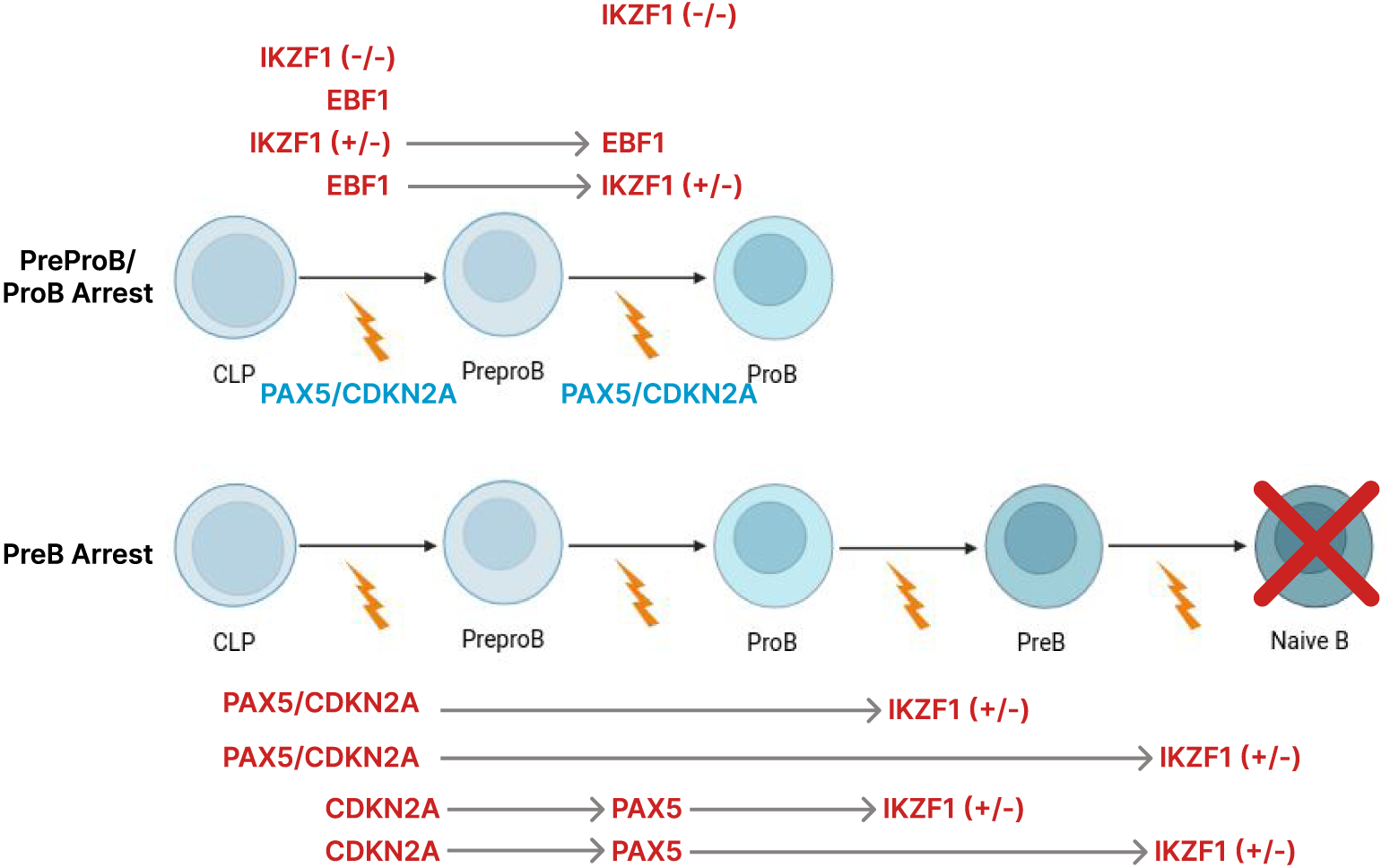
Mapping the Hierarchical Evolutionary Trajectories of *BCR-ABL* BCP-ALL. Schematic representation of the most probable genetic paths driving leukemogenesis, identified through computational filtering for developmental reachability and stepwise fitness optimization. **Top Panel**: Evolutionary Trajectories resulting in early maturation arrest. These trajectories originate at the CLP stage and culminate in a stable differentiation block at the Pre-pro-B or Pro-B stages. These paths are primarily driven by early-window hits to *IKZF1* (both -/- and +/-) or *EBF1* (+/-). Notably, these evolutionary trajectories accommodate clinical profiles lacking *PAX5* or *CDKN2A* alterations, demonstrating that early-stage blocks can occur independently of downstream perturbations. **Bottom Panel**: Evolutionary Trajectories resulting in late maturation arrest. These paths illustrate the progression toward a Pre-B arrest. The lightning bolt symbols indicate points of genetic alteration used in the simulation to perturb core regulatory nodes.

## 4 Discussion

Here we present the first computational model of B-cell development using multi-valued logic, centred around a core network of transcription factors - themselves frequent targets for mutation in BCP-ALL. We used this as a framework to model leukaemogenesis, taking *BCR-ABL1* as a subtype-defining initiating event. The model allows us to explore alternative evolutionary trajectories where mutations are acquired at different developmental time points, and in different combinations. This reveals that the point of maturation arrest and the fitness of the end-state are dependent not only on the particular combination of mutations but on the order in which they are acquired. We therefore propose that the different states of *BCR-ABL1* BCP-ALL observed among patients [16] reflect alternative evolutionary trajectories with implications for targeted therapy and interception strategies.

The complex regulatory landscape of BCP-ALL necessitates a modelling approach that differs from Boolean formalism frequently used in computational cancer biology. We selected a multi-valued logical formalism to explicitly represent non-binary, graded activity levels. Unlike standard Boolean models, this approach captures dosage effects—specifically distinguishing between homozygous and heterozygous losses—which is fundamental to modelling the impacts of mutations in the core transcription factors on B-cell maturation. While healthy B-cell development may be modelled as a fixed sequential program, the genetic alterations within this framework are not temporally restricted; represented in modelling formalism as deterministic and stochastic processes respectively. We resolved this by introducing genetic alterations at varying time points within deterministic simulations, effectively recapitulating the stochastic nature of mutation in leukaemogenesis within a stable computational framework. The integration of *BCR-ABL1* signalling into the B-cell GRN scaffold, and the modelling of diverse classes of mutations, represents the first computational effort to describe the mechanistic trajectories from healthy maturation to leukaemic arrest. This establishes a predictive framework for determining how the timing and order of genetic insults dictate the maturation arrest point and the fitness of the leukaemic clone. The treatment of concurrency in our model is more nuanced than the commonly chosen synchronous/deterministic or asynchronous/stochastic updating scheme. It is increasingly recognized that different biological systems need sophisticated and bespoke concurrency schemes [47, 48]. Our work reinforces this conclusion, and leukamogenesis gives an example of the specific and widespread phenomena of fixed and sequential developmental systems that can accrue mutations at arbitrary times. This could include other paediatric cancers, but also potentially the influence of epigenetic clocks in aging.

Our simulations uncover the internal regulatory rules that determine which specific leukaemic genotypes are observed in tumours within the *BCR-ABL1* subtype. A critical insight from our trajectory analysis is that the developmental window in which an alteration occurs is as decisive as the alteration itself. For instance, the early loss of *IKZF1* or *EBF1* effectively traps the cell in a Pre-pro-B state, rendering subsequent alterations that might otherwise drive a later Pre-B arrest biologically unreachable. Conversely, loss of *IKZF1* beyond the proB stage cannot effect differentiation arrest but still impacts fitness. Our unbiased simulation of 784 permutations across 17 clinically-observed genetic profiles identified that only 147 paths are biologically feasible. Filtering for trajectories in which each additional mutation must provide a selective advantage, we narrowed the results to a core set of 58 highly probable evolutionary paths. This suggested that rather than evolving through a “random” accumulation of genetic alterations, BCP-ALL arises in the context of a highly constrained evolutionary landscape.

The model also allowed us to explore the different end-states arising from alternative evolutionary trajectories. We observed a fundamental distinction between fixed-point attractors which represent a stable phenotype, and cyclic attractors which are characterized by perpetual phenotypic fluctuation and particularly associated with combined *EBF1* and *IKZF1* heterozygous loss. Cyclic attractors potentially signify a state of continuous regulatory flux cycles of attempted differentiation and signalling-induced stress. These internal dynamics increase the diversity of states in which additional mutations can have an impact and could explain why certain molecular profiles are predisposed to bypass cellular monitoring. These features of our model highlight its ability to shed light on clinically silent phases of leukaemogenesis that may determine heterogeneity observed between patients.

We have constructed a model of B-cell differentiation that spans the developmental stages in which differentiation arrest occurs in BCP-ALL. This has allowed us to explore the impact of different orders and timing of mutations. However, the use of a limited number of markers may excessively discretize what is biologically a maturation continuum. We envisage that future iterations of the model will include additional markers, such as the canonical cell surface markers *CD34*, *CD19*, and *CD10* [49], to describe cell states with increased granularity. Furthermore, while we have successfully modelled loss-of-function mutations, increasing the range of activity levels within the multi-valued logic framework would allow us to model gain-of-function. While we have chosen to focus on BCR-ABL and the most frequent second hits in this subtype there are many alternative initiating events and secondary mutations that can combine to drive BCP-ALL [7]. Our model can readily be adapted to study these, providing an opportunity to uncover the shared and divergent regulatory logics that govern leukaemic transformation across the entire B-cell precursor landscape.

Our approach not only distinguishes alternative end-states but provides a method to distinguish the most probable evolutionary trajectories among many possibilities by comparing the fitness of intermediate steps. This provides a window into the pre-malignant phase of the disease - not normally observed clinically - and may reveal opportunities for disease interception. Furthermore, the model could be used to simulate the impact of targeted treatments across genetic profiles, potentially informing a more personalized approach to treatment.

This platform provides a generalizable framework for predicting evolutionary trajectories in multi-hit malignancies where there is sufficient knowledge of the mutational spectrum and cellular differentiation hierarchies.

## 5 Acknowledgments

We thank CRUK for funding this project (grant reference CDEPIL-Jan24/100032). TE acknowledges support from Blood Cancer UK (grant references 21003 and 24017). We thank Gill May and Virginia Turati for their help in the review of this manuscript.

## 6 Author contributions

MB built the model, performed all calculations, and wrote the initial drafts of the manuscript. BH wrote software used in the study. JW supported model development and literature curation. BH and TE designed and supervised the study. TE and BH contributed equally to this work. All authors edited, reviewed, and contributed to writing the submitted manuscript.

## Supplementary information

**SFig 1**: **Stability analysis of the model in the CLP state in the absence of exogenous inputs *IL7*, *IKZF1*).**

**SFig 2**: **Stability analysis of the model in the Naive B-cell state.**

**SFig 3**: **BMA model map.**

**SFig 4**: **Long Sensitivity analysis of the model following individual TF alterations.**

**SFig 5**: **Characterization of filtered secondary hit hierarchies and evolutionary trajectories in *BCR-ABL* BCP-ALL.**

**SFig 6**: **Characterization of Filtered secondary hit hierarchies and evolutionary trajectories in *BCR-ABL* BCP-ALL.**

**SFig 7: Characterization of filtered secondary hit hierarchies and evolutionary trajectories in *BCR-ABL* BCP-ALL compared to *BCR-ABL* only. A–C.**

**SFig 8: Heatmap of stage-specific marker dynamics under *BCR-ABL* and *EBF1* second hit conditions.**

**SFig 9: Heatmap of stage-specific marker dynamics under *BCR-ABL* and *EBF1* second hit conditions.**

**SFig 10: Heatmap of stage-specific marker dynamics under *BCR-ABL* and *IKZF1* second hit conditions.**

**SFig 11: Heatmap of stage-specific marker dynamics under *BCR-ABL* and *PAX5* second hit conditions.**

**SFig 12: Heatmap of 58 “highly probable” trajectories.**

**STable 1**: **Model interaction table**

## Footnotes

1 https://github.com/hallba/pybma/tree/main

## Notes

https://doi.org/10.5281/zenodo.20135289

### Competing Interest Statement

The authors have declared no competing interest.

## References

[1] Inaba, H., Mullighan, C.G.: Pediatric acute lymphoblastic leukemia. Haematologica 105(11), 2524 (2020)

[2] Pui, C.-H., Carroll, W.L., Meshinchi, S., Arceci, R.J.: Biology, risk stratification, and therapy of pediatric acute leukemias: an update. Journal of clinical oncology 29(5), 551–565 (2011)

[3] Greaves, M.: A causal mechanism for childhood acute lymphoblastic leukaemia. Nature Reviews Cancer 18(8), 471–484 (2018)

[4] Weyden, L., Giotopoulos, G., Rust, A.G., Matheson, L.S., Delft, F.W., Kong, J., Corcoran, A.E., Greaves, M.F., Mullighan, C.G., Huntly, B.J., et al.: Modeling the evolution of etv6-runx1–induced b-cell precursor acute lymphoblastic leukemia in mice. Blood, The Journal of the American Society of Hematology 118(4), 1041–1051 (2011)

[5] Fischer, U., Yang, J.J., Ikawa, T., Hein, D., Vicente-Dueñas, C., Borkhardt, A., Sánchez-García, I.: Cell fate decisions: the role of transcription factors in early b-cell development and leukemia. Blood cancer discovery 1(3), 224–233 (2020)

[6] Schwab, C.J., Chilton, L., Morrison, H., Jones, L., Al-Shehhi, H., Erhorn, A., Russell, L.J., Moorman, A.V., Harrison, C.J.: Genes commonly deleted in childhood b-cell precursor acute lymphoblastic leukemia: association with cytogenetics and clinical features. Haematologica 98(7), 1081 (2013)

[7] Brady, S.W., Roberts, K.G., Gu, Z., Shi, L., Pounds, S., Pei, D., Cheng, C., Dai, Y., Devidas, M., Qu, C., et al.: The genomic landscape of pediatric acute lymphoblastic leukemia. Nature genetics 54(9), 1376–1389 (2022)

[8] Hunger, S.P., Mullighan, C.G.: Redefining all classification: toward detecting high-risk all and implementing precision medicine. Blood, The Journal of the American Society of Hematology 125(26), 3977–3987 (2015)

[9] Collombet, S., Van Oevelen, C., Sardina Ortega, J.L., Abou-Jaoudé, W., Di Stefano, B., Thomas-Chollier, M., Graf, T., Thieffry, D.: Logical modeling of lymphoid and myeloid cell specification and transdifferentiation. Proceedings of the National Academy of Sciences 114(23), 5792–5799 (2017)

[10] Mendoza, L., Méndez, A.: A dynamical model of the regulatory network controlling lymphopoiesis. Biosystems 137, 26–33 (2015)

[11] Méndez, A., Mendoza, L.: A network model to describe the terminal differentiation of b cells. PLoS computational biology 12(1), 1004696 (2016)

[12] Benque, D., Bourton, S., Cockerton, C., Cook, B., Fisher, J., Ishtiaq, S., Piterman, N., Taylor, A., Vardi, M.Y.: Bma: Visual tool for modeling and analyzing biological networks. In: International Conference on Computer Aided Verification, pp. 686–692 (2012). Springer

[13] Campos-Sanchez, E., Toboso-Navasa, A., Romero-Camarero, I., Barajas-Diego, M., Sanchez-Garcia, I., Cobaleda, C.: Acute lymphoblastic leukemia and developmental biology: a crucial interrelationship. Cell Cycle 10(20), 3473–3486 (2011)

[14] Lee, R.D., Munro, S.A., Knutson, T.P., LaRue, R.S., Heltemes-Harris, L.M., Farrar, M.A.: Single-cell analysis identifies dynamic gene expression networks that govern b cell development and transformation. Nature communications 12(1), 6843 (2021)

[15] Pieper, K., Grimbacher, B., Eibel, H.: B-cell biology and development. Journal of Allergy and Clinical Immunology 131(4), 959–971 (2013)

[16] Kim, J.C., Chan-Seng-Yue, M., Ge, S., Zeng, A.G., Ng, K., Gan, O.I., Garcia-Prat, L., Flores-Figueroa, E., Woo, T., Zhang, A.X.W., et al.: Transcriptomic classes of bcr-abl1 lymphoblastic leukemia. Nature genetics 55(7), 1186–1197 (2023)

[17] Sigvardsson, M.: Transcription factor networks link blymphocyte development and malignant transformation in leukemia. Genes & Development 37(15-16), 703–723 (2023)

[18] Kanehisa, M., Goto, S.: Kegg: kyoto encyclopedia of genes and genomes. Nucleic acids research 28(1), 27–30 (2000)

[19] Fedl, A.S., Tagoh, H., Gruenbacher, S., Sun, Q., Schenk, R.L., Froussios, K., Jaritz, M., Busslinger, M., Schwickert, T.A.: Transcriptional function of e2a, ebf1, pax5, ikaros and aiolos analyzed by in vivo acute protein degradation in early b cell development. Nature Immunology 25(9), 1663–1677 (2024)

[20] Ferreirós-Vidal, I., Carroll, T., Taylor, B., Terry, A., Liang, Z., Bruno, L., Dharmalingam, G., Khadayate, S., Cobb, B.S., Smale, S.T., et al.: Genome-wide identification of ikaros targets elucidates its contribution to mouse b-cell lineage specification and pre-b–cell differentiation. Blood, The Journal of the American Society of Hematology 121(10), 1769–1782 (2013)

[21] Kee, B.L., Murre, C.: Induction of early b cell factor (ebf) and multiple b lineage genes by the basic helix-loop-helix transcription factor e12. The Journal of experimental medicine 188(4), 699–713 (1998)

[22] Kikuchi, K., Lai, A.Y., Hsu, C.-L., Kondo, M.: Il-7 receptor signaling is necessary for stage transition in adult b cell development through up-regulation of ebf. The Journal of experimental medicine 201(8), 1197–1203 (2005)

[23] Zandi, S., Mansson, R., Tsapogas, P., Zetterblad, J., Bryder, D., Sigvardsson, M.: Ebf1 is essential for b-lineage priming and establishment of a transcription factor network in common lymphoid progenitors. The Journal of Immunology 181(5), 3364–3372 (2008)

[24] Lin, H., Grosschedl, R.: Failure of b-cell differentiation in mice lacking the transcription factor ebf. Nature 376(6537), 263–267 (1995)

[25] Heavey, B., Charalambous, C., Cobaleda, C., Busslinger, M.: Myeloid lineage switch of pax5 mutant but not wild-type b cell progenitors by c/ebp*α* and gata factors. The EMBO Journal (2003)

[26] O’Riordan, M., Grosschedl, R.: Coordinate regulation of b cell differentiation by the transcription factors ebf and e2a. Immunity 11(1), 21–31 (1999)

[27] Li, L., Zhang, D., Cao, X.: Ebf1, pax5, and myc: regulation on b cell development and association with hematologic neoplasms. Frontiers in Immunology 15, 1320689 (2024)

[28] García-Gutiérrez, L., Bretones, G., Molina, E., Arechaga, I., Symonds, C., Acosta, J.C., Blanco, R., Fernández, A., Alonso, L., Sicinski, P., et al.: Myc stimulates cell cycle progression through the activation of cdk1 and phosphorylation of p27. Scientific reports 9(1), 18693 (2019)

[29] Geng, H., Hurtz, C., Lenz, K.B., Chen, Z., Baumjohann, D., Thompson, S., Goloviznina, N.A., Chen, W.-Y., Huan, J., LaTocha, D., et al.: Self-enforcing feed-back activation between bcl6 and pre-b cell receptor signaling defines a distinct subtype of acute lymphoblastic leukemia. Cancer cell 27(3), 409–425 (2015)

[30] Swaminathan, S., Huang, C., Geng, H., Chen, Z., Harvey, R., Kang, H., Ng, C., Titz, B., Hurtz, C., Sadiyah, M.F., et al.: Bach2 mediates negative selection and p53-dependent tumor suppression at the pre-b cell receptor checkpoint. Nature medicine 19(8), 1014–1022 (2013)

[31] Bain, G., Maandag, E.C.R., Izon, D.J., Amsen, D., Kruisbeek, A.M., Weintraub, B.C., Krop, I., Schlissel, M.S., Feeney, A.J., Roon, M., et al.: E2a proteins are required for proper b cell development and initiation of immunoglobulin gene rearrangements. Cell 79(5), 885–892 (1994)

[32] Nechanitzky, R., Akbas, D., Scherer, S., Györy, I., Hoyler, T., Ramamoorthy, S., Diefenbach, A., Grosschedl, R.: Transcription factor ebf1 is essential for the maintenance of b cell identity and prevention of alternative fates in committed cells. Nature immunology 14(8), 867–875 (2013)

[33] Mansson, R., Welinder, E., Åhsberg, J., Lin, Y.C., Benner, C., Glass, C.K., Lucas, J.S., Sigvardsson, M., Murre, C.: Positive intergenic feedback circuitry, involving ebf1 and foxo1, orchestrates b-cell fate. Proceedings of the National Academy of Sciences 109(51), 21028–21033 (2012)

[34] Urbánek, P., Wang, Z.-Q., Fetka, I., Wagner, E.F., Busslinger, M.: Complete block of early b cell differentiation and altered patterning of the posterior midbrain in mice lacking pax5bsap. Cell 79(5), 901–912 (1994)

[35] Hu, Y., Yoshida, T., Georgopoulos, K.: Transcriptional circuits in b cell transformation. Current opinion in hematology 24(4), 345–352 (2017)

[36] Yoshida, T., Yao-Ming Ng, S., Zuniga-Pflucker, J.C., Georgopoulos, K.: Early hematopoietic lineage restrictions directed by ikaros. Nature immunology 7(4), 382–391 (2006)

[37] Ng, S.Y.-M., Yoshida, T., Zhang, J., Georgopoulos, K.: Genome-wide lineage-specific transcriptional networks underscore ikaros-dependent lymphoid priming in hematopoietic stem cells. Immunity 30(4), 493–507 (2009)

[38] Georgopoulos, K., Bigby, M., Wang, J.-H., Molnar, A., Wu, P., Winandy, S., Sharpe, A.: The ikaros gene is required for the development of all lymphoid lineages. Cell 79(1), 143–156 (1994)

[39] Dengler, H.S., Baracho, G.V., Omori, S.A., Bruckner, S., Arden, K.C., Castrillon, D.H., DePinho, R.A., Rickert, R.C.: Distinct functions for the transcription factor foxo1 at various stages of b cell differentiation. Nature immunology 9(12), 1388–1398 (2008)

[40] Duy, C., Hurtz, C., Shojaee, S., Cerchietti, L., Geng, H., Swaminathan, S., Klemm, L., Kweon, S.-m., Nahar, R., Braig, M., et al.: Bcl6 enables ph+ acute lymphoblastic leukaemia cells to survive bcr–abl1 kinase inhibition. Nature 473(7347), 384–388 (2011)

[41] Casolari, D., Makri, M., Yoshida, C., Muto, A., Igarashi, K., Melo, J.: Transcriptional suppression of bach2 by the bcrabl oncoprotein is mediated by pax5. Leukemia 27(2), 409–415 (2013)

[42] Chuang, R., Hall, B.A., Benque, D., Cook, B., Ishtiaq, S., Piterman, N., Taylor, A., Vardi, M., Koschmieder, S., Gottgens, B., et al.: Drug target optimization in chronic myeloid leukemia using innovative computational platform. Scientific reports 5(1), 8190 (2015)

[43] Iacobucci, I., Zeng, A.G., Gao, Q., Garcia-Prat, L., Baviskar, P., Shah, S., Muri-son, A., Voisin, V., Chan-Seng-Yue, M., Cheng, C., et al.: Multipotent lineage potential in b cell acute lymphoblastic leukemia is associated with distinct cellular origins and clinical features. Nature Cancer, 1–26 (2025)

[44] Mullighan, C.G., Miller, C.B., Radtke, I., Phillips, L.A., Dalton, J., Ma, J., White, D., Hughes, T.P., Le Beau, M.M., Pui, C.-H., et al.: Bcr–abl1 lymphoblastic leukaemia is characterized by the deletion of ikaros. Nature 453(7191), 110–114 (2008)

[45] Chang, T.-C., Chen, W., Qu, C., Cheng, Z., Hedges, D., Elsayed, A., Pounds, S.B., Shago, M., Rabin, K.R., Raetz, E.A., et al.: Genomic determinants of outcome in acute lymphoblastic leukemia. Journal of Clinical Oncology 42(29), 3491–3503 (2024)

[46] Mullighan, C.G., Su, X., Zhang, J., Radtke, I., Phillips, L.A., Miller, C.B., Ma, J., Liu, W., Cheng, C., Schulman, B.A., et al.: Deletion of ikzf1 and prognosis in acute lymphoblastic leukemia. New England Journal of Medicine 360(5), 470–480 (2009)

[47] Paulevé, L., Kolčák, J., Chatain, T., Haar, S.: Reconciling qualitative, abstract, and scalable modeling of biological networks. Nature communications 11(1) (2020)

[48] Lim, C.Y., Wang, H., Woodhouse, S., Piterman, N., Wernisch, L., Fisher, J., Göttgens, B.: Btr: training asynchronous boolean models using single-cell expression data. BMC bioinformatics, 17(1) (2016)

[49] Van Lochem, E., Velden, V., Wind, H., Te Marvelde, J., Westerdaal, N., Van Dongen, J.: Immunophenotypic differentiation patterns of normal hematopoiesis in human bone marrow: reference patterns for age-related changes and disease-induced shifts. Cytometry Part B: Clinical Cytometry: The Journal of the International Society for Analytical Cytology 60(1), 1–13 (2004)

