## Supplementary figures and images for "Dynamics of the B-Cell gene regulatory network in differentiation determine evolutionary trajectories of childhood leukaemogenesis"

### Fig1.pdf

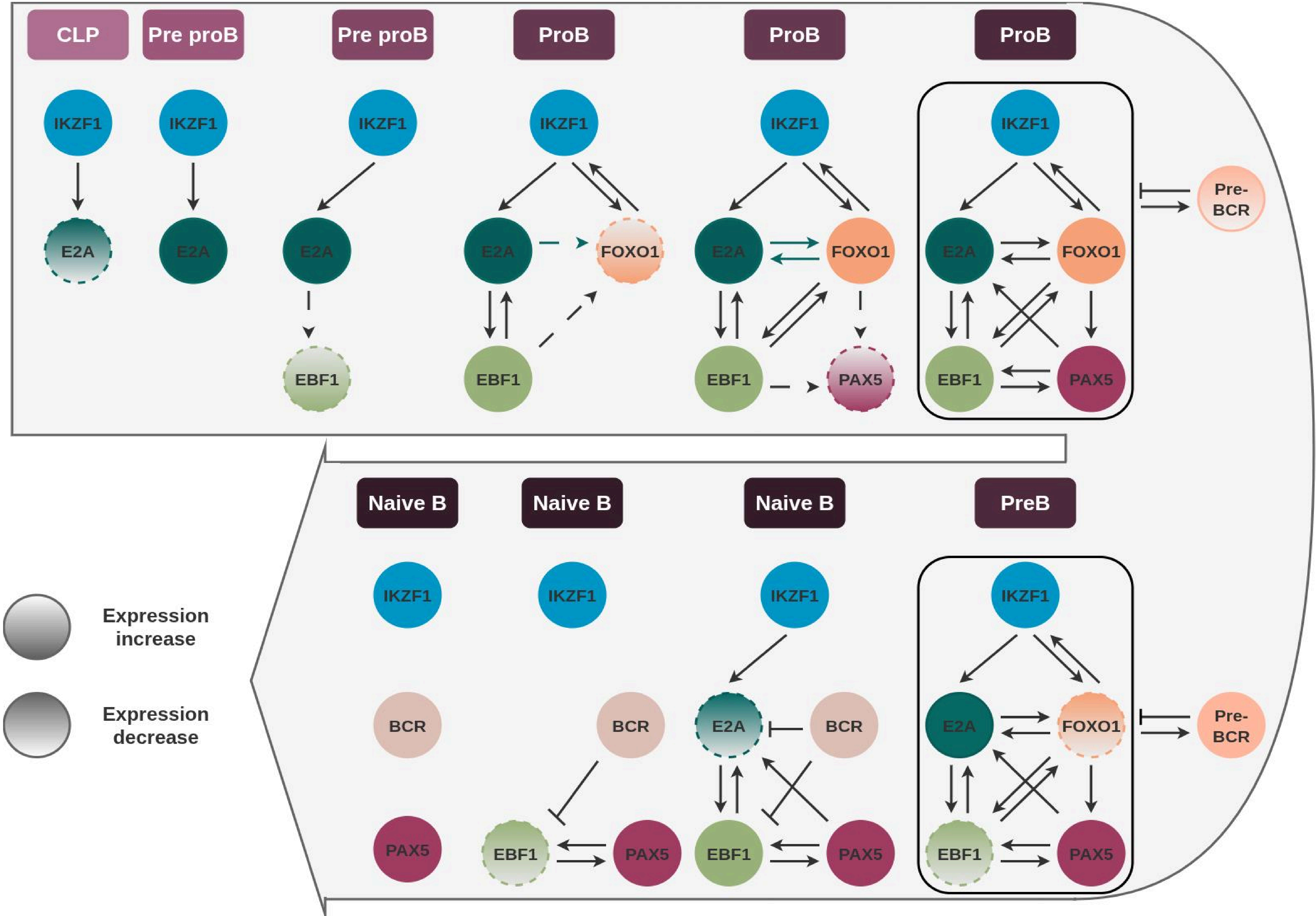

### Fig2.pdf

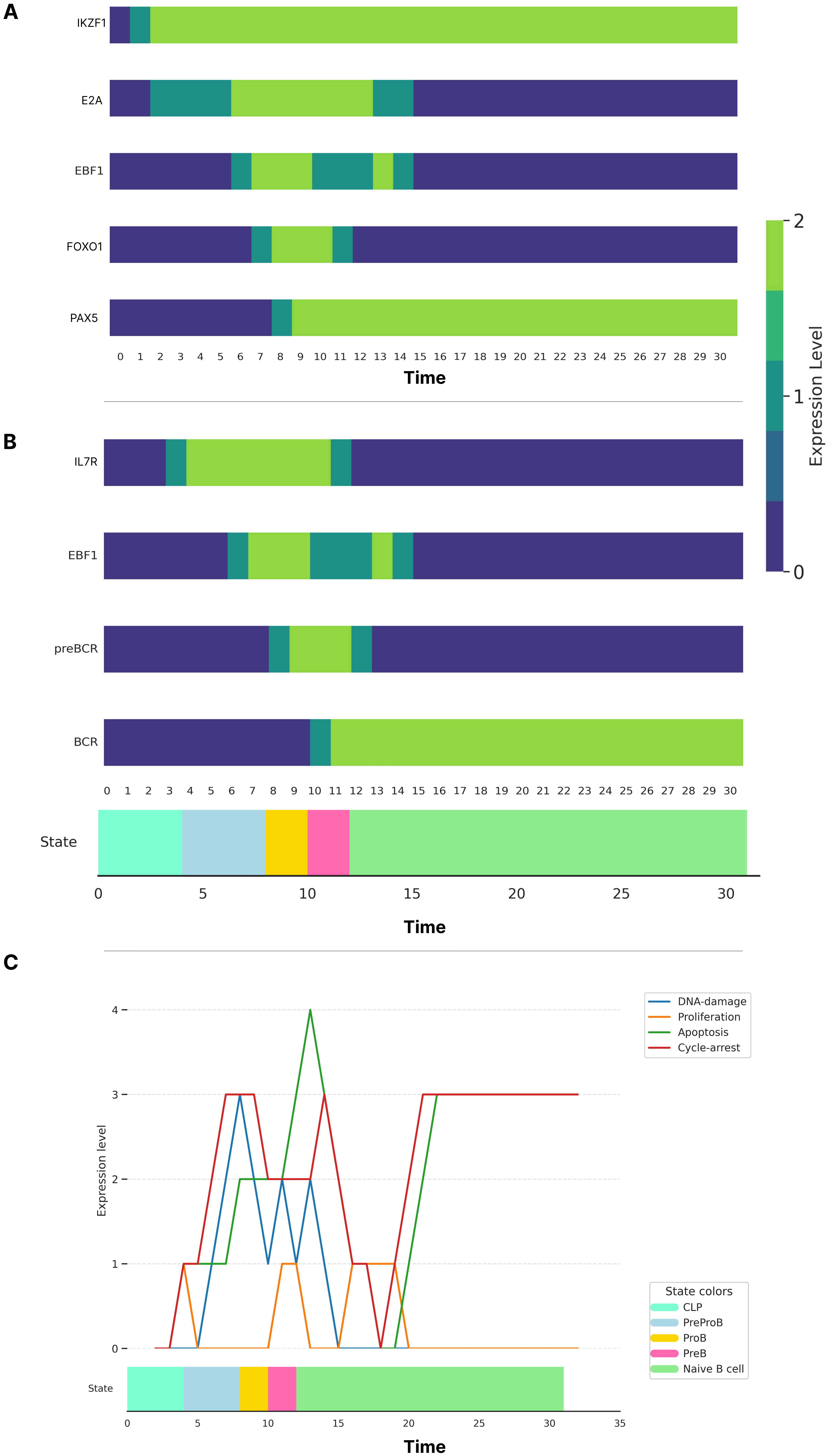

### Fig3.pdf

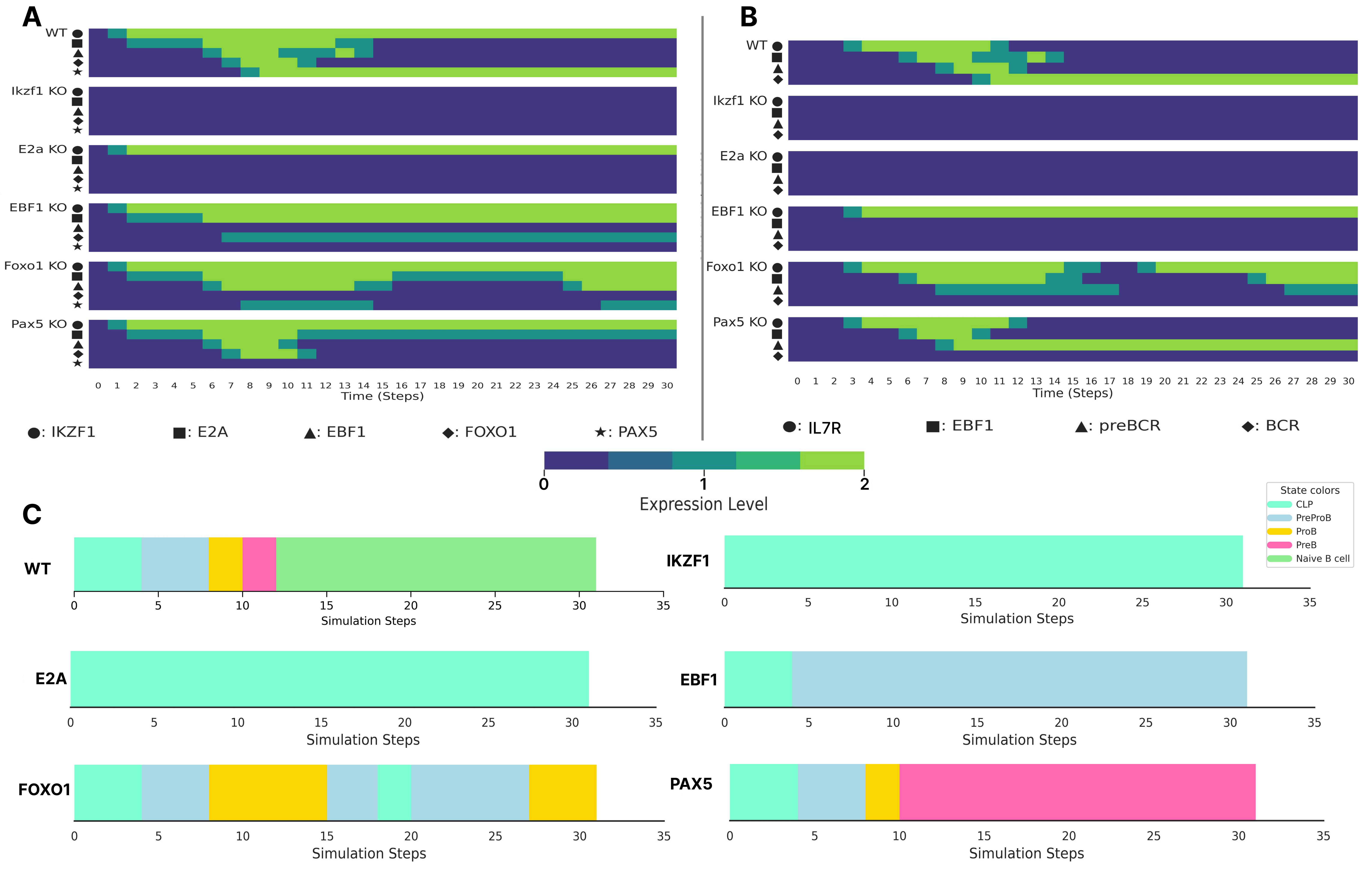

### Fig4.pdf

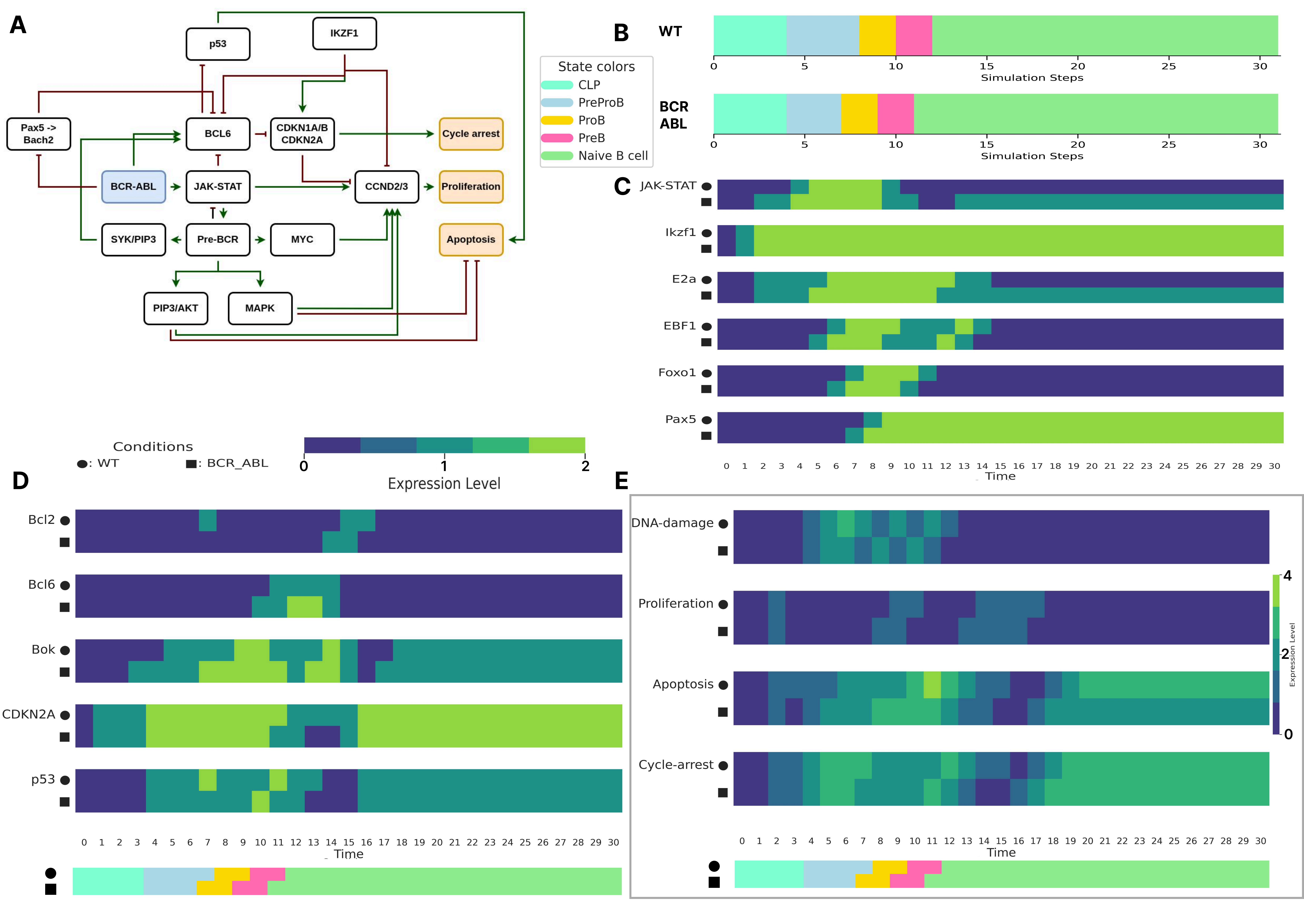

### Fig5.pdf

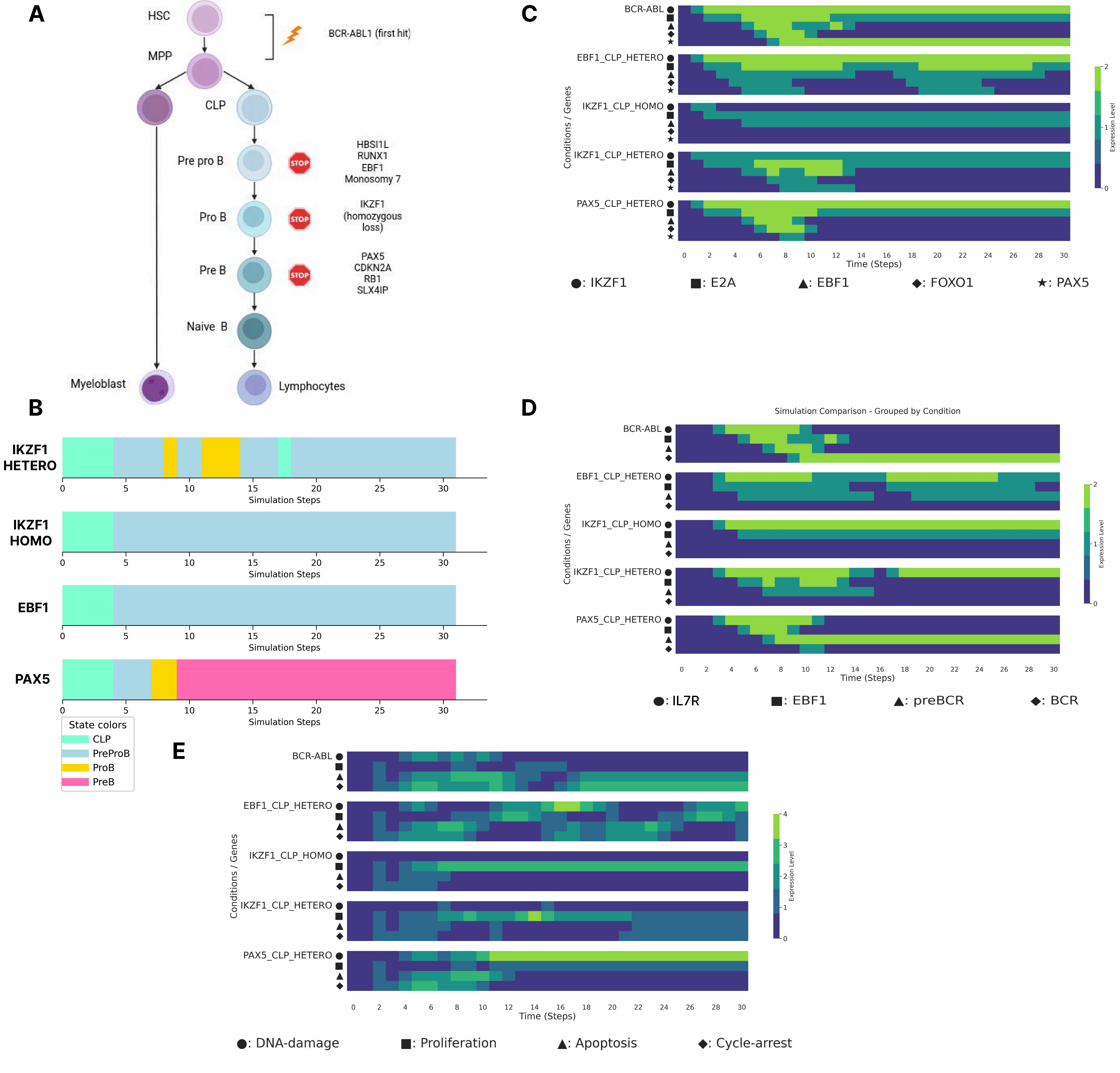

### Fig6.pdf

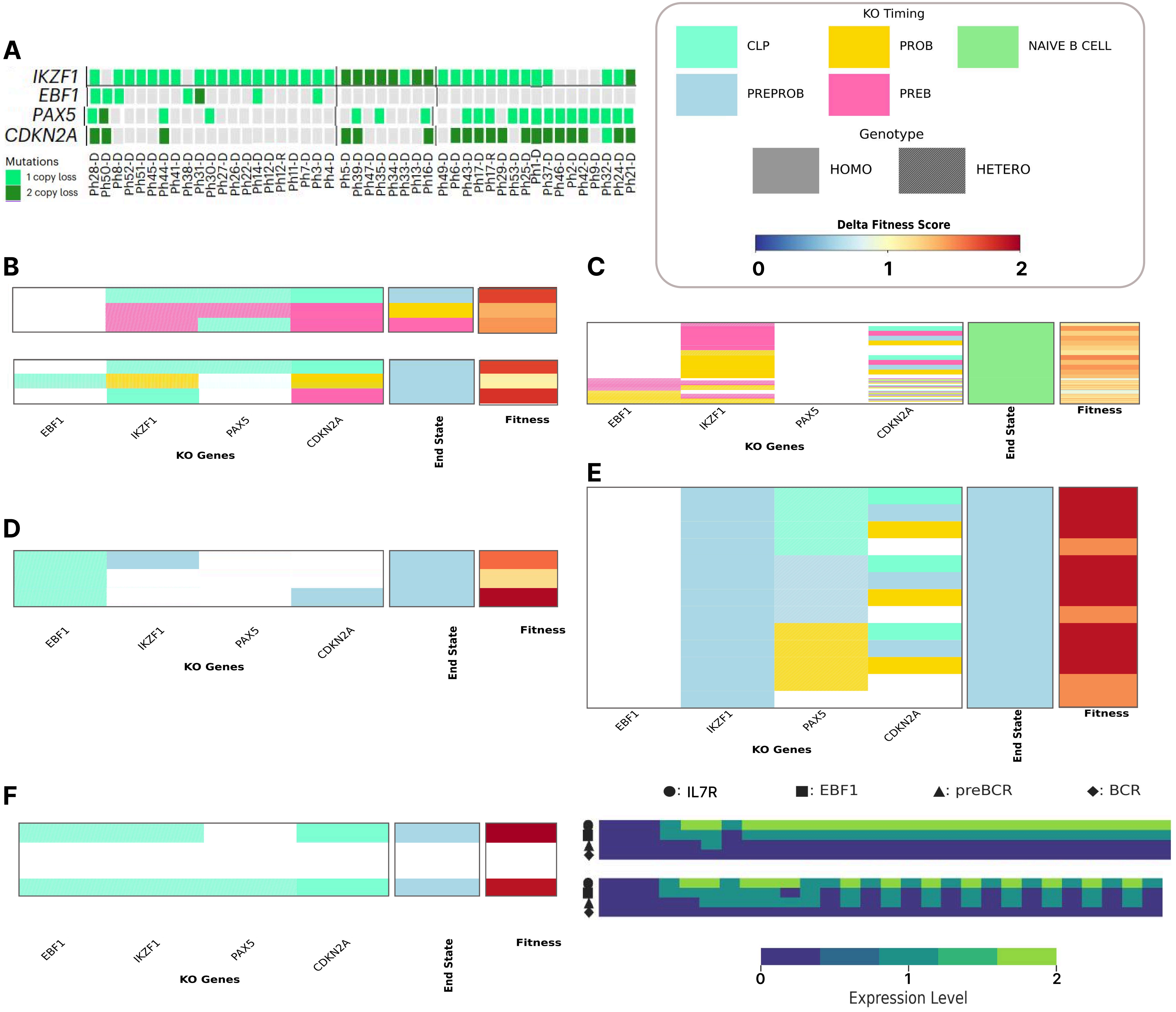

### Fig7.pdf

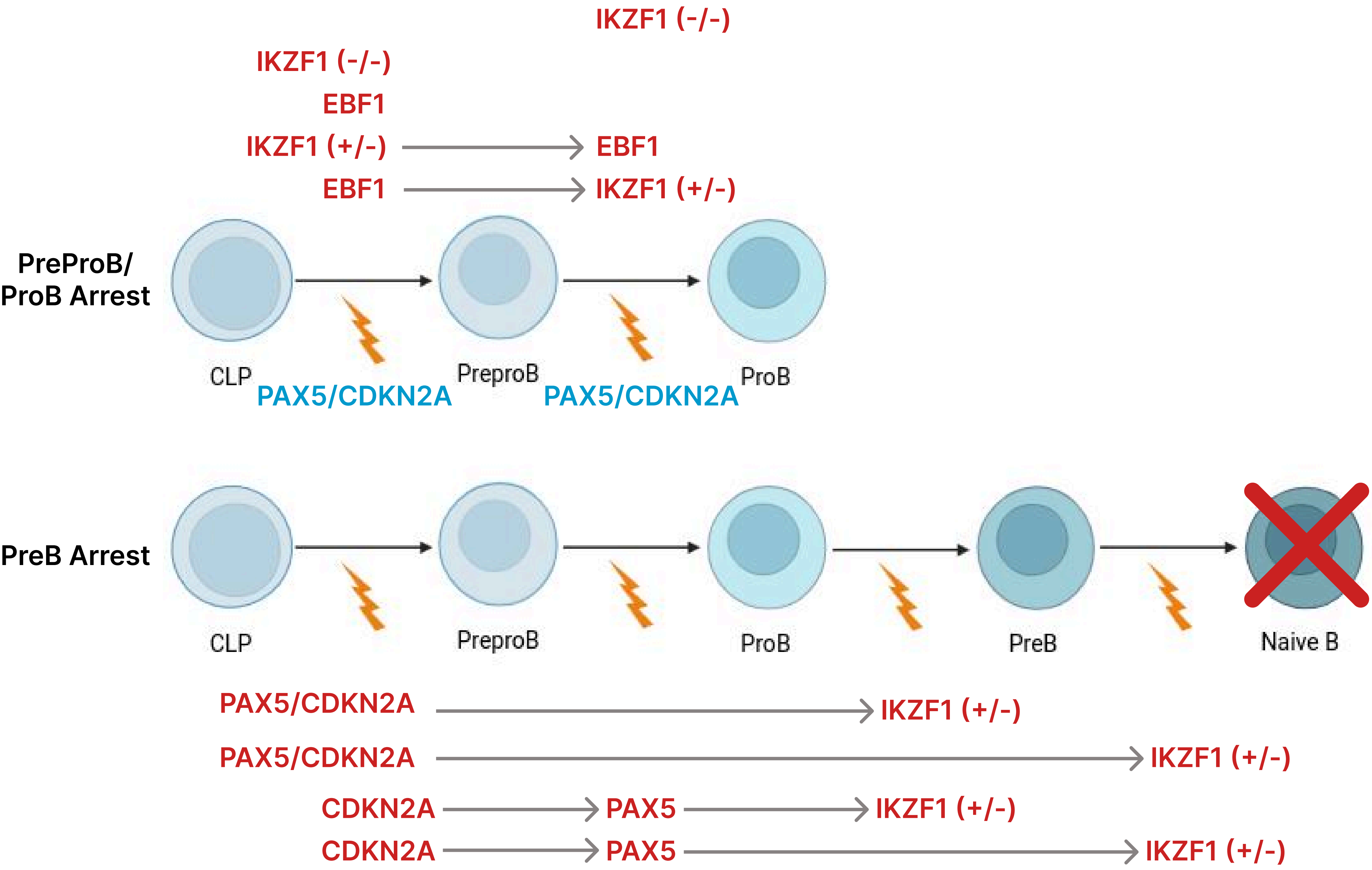

### S1-WT-CLP-stability-analysis.png

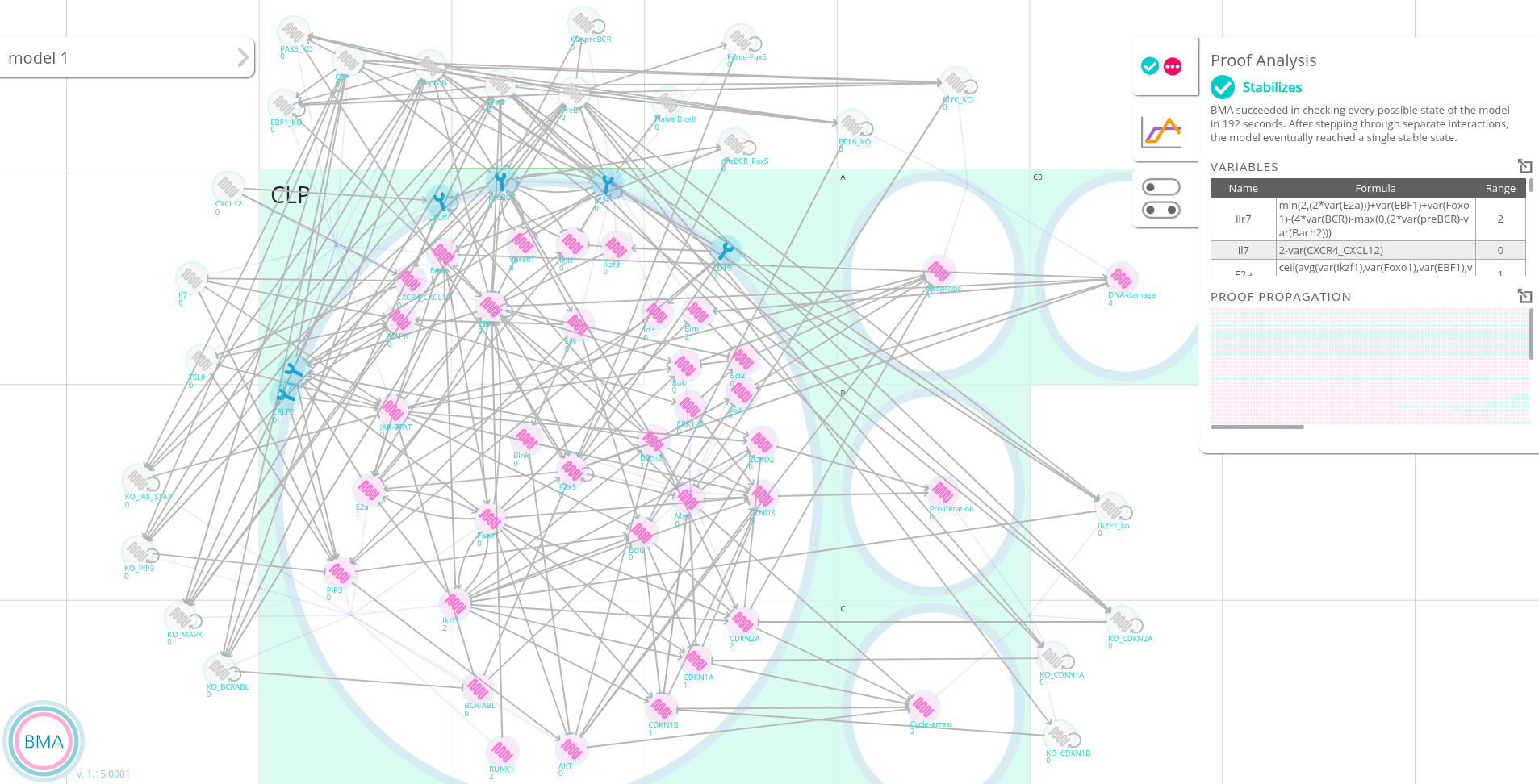

### S2-WT-NaiveB-stability-analysis.png

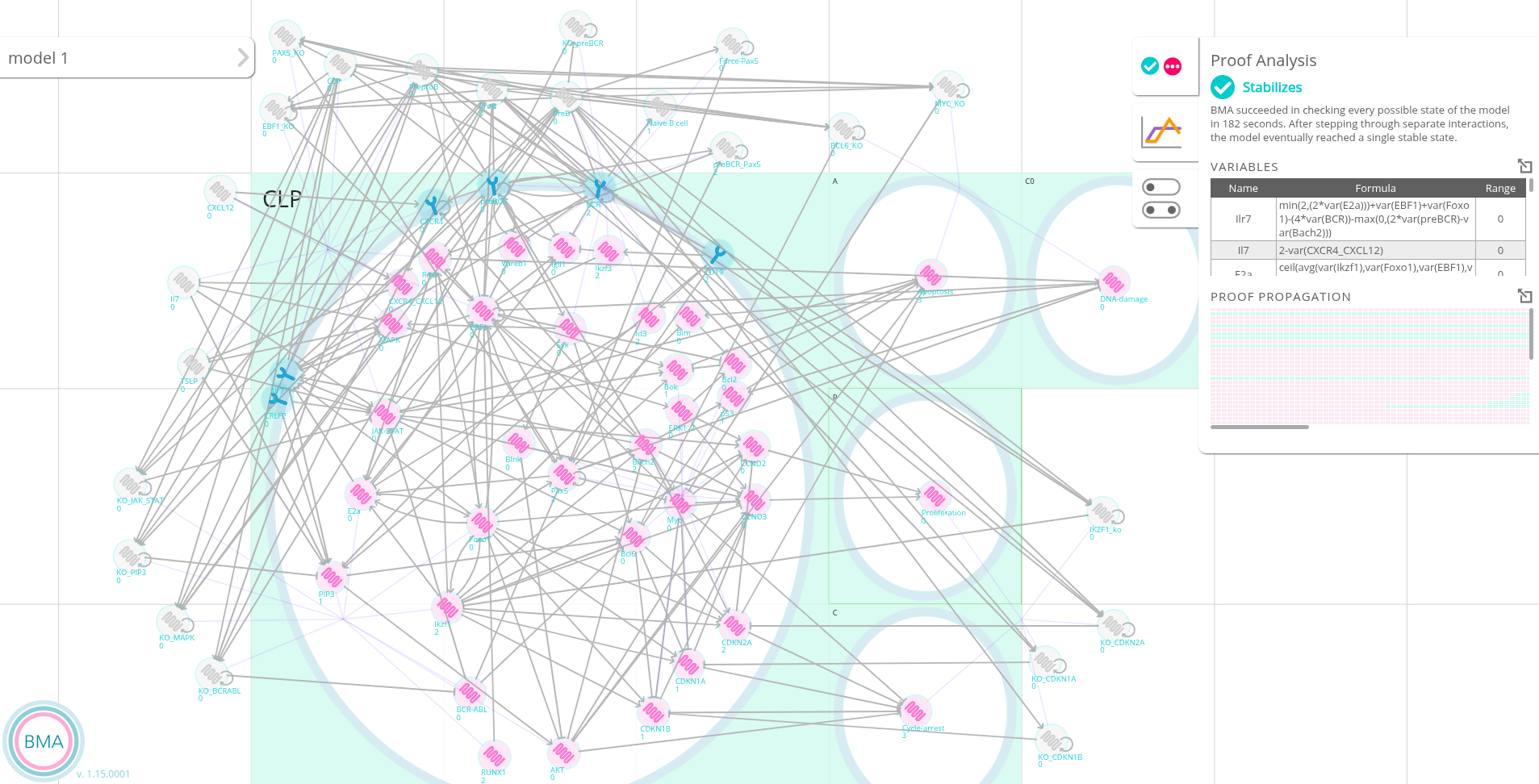

### S3Fig.pdf

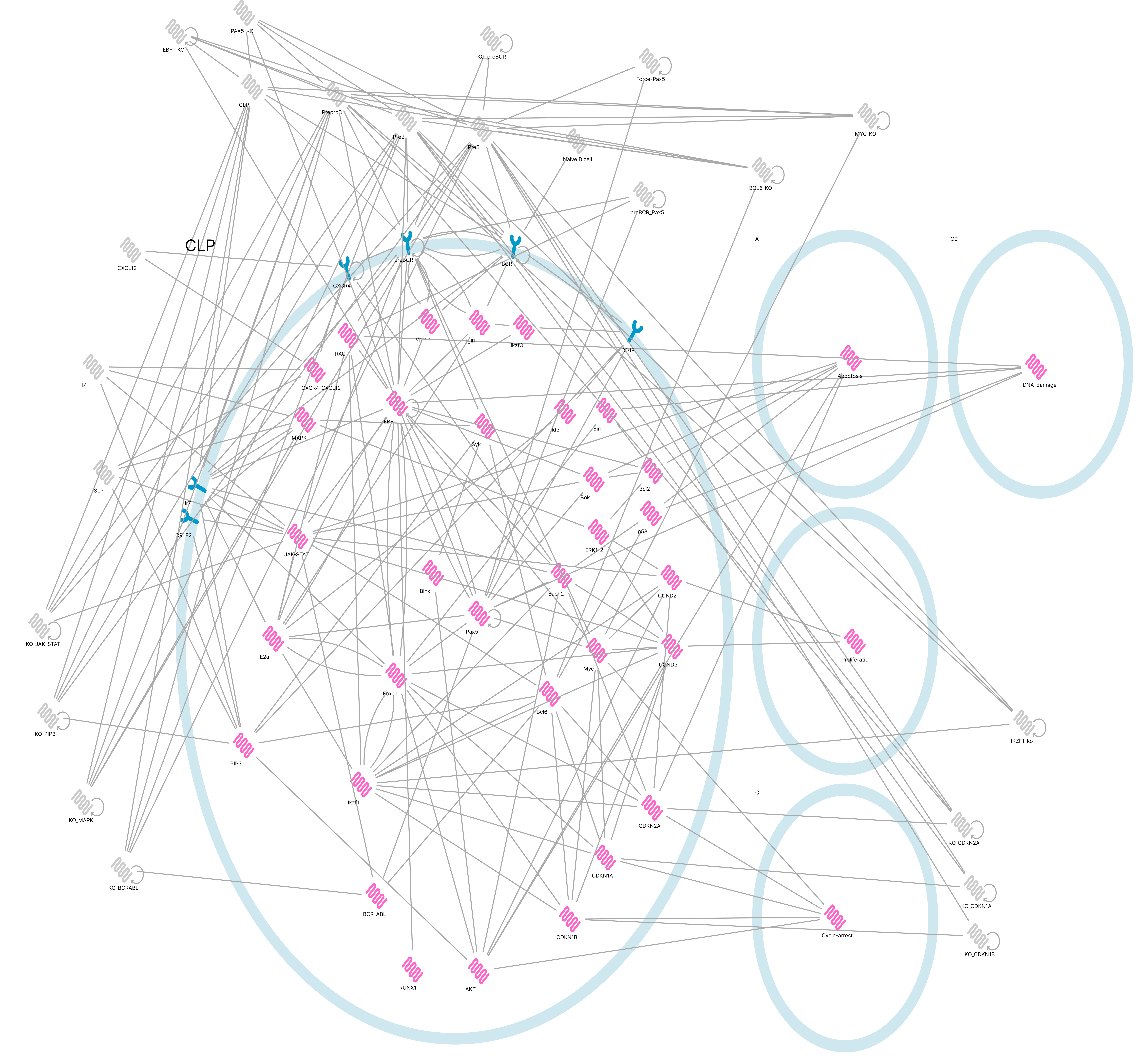

### S4Fig.pdf

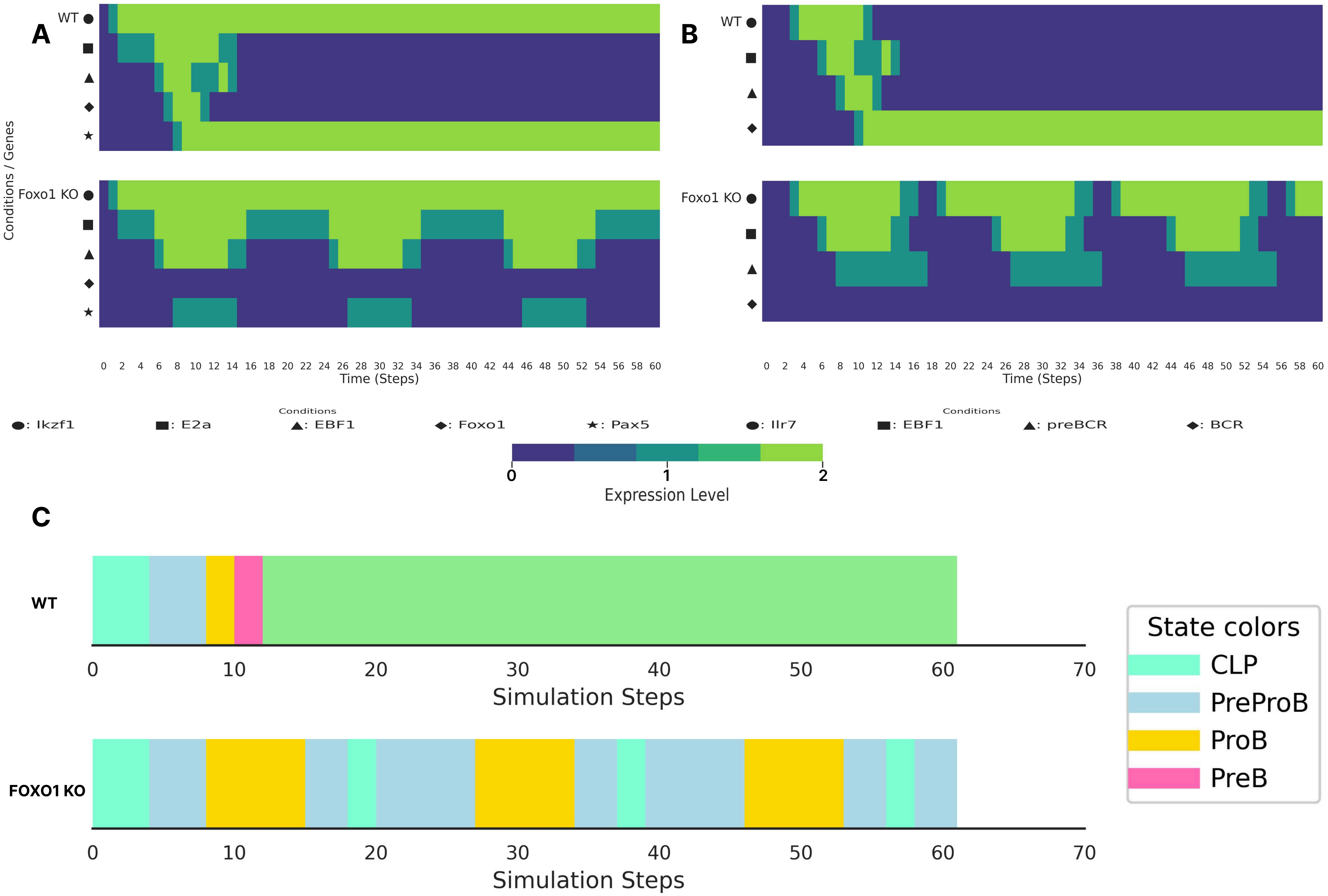

### S5Fig.pdf

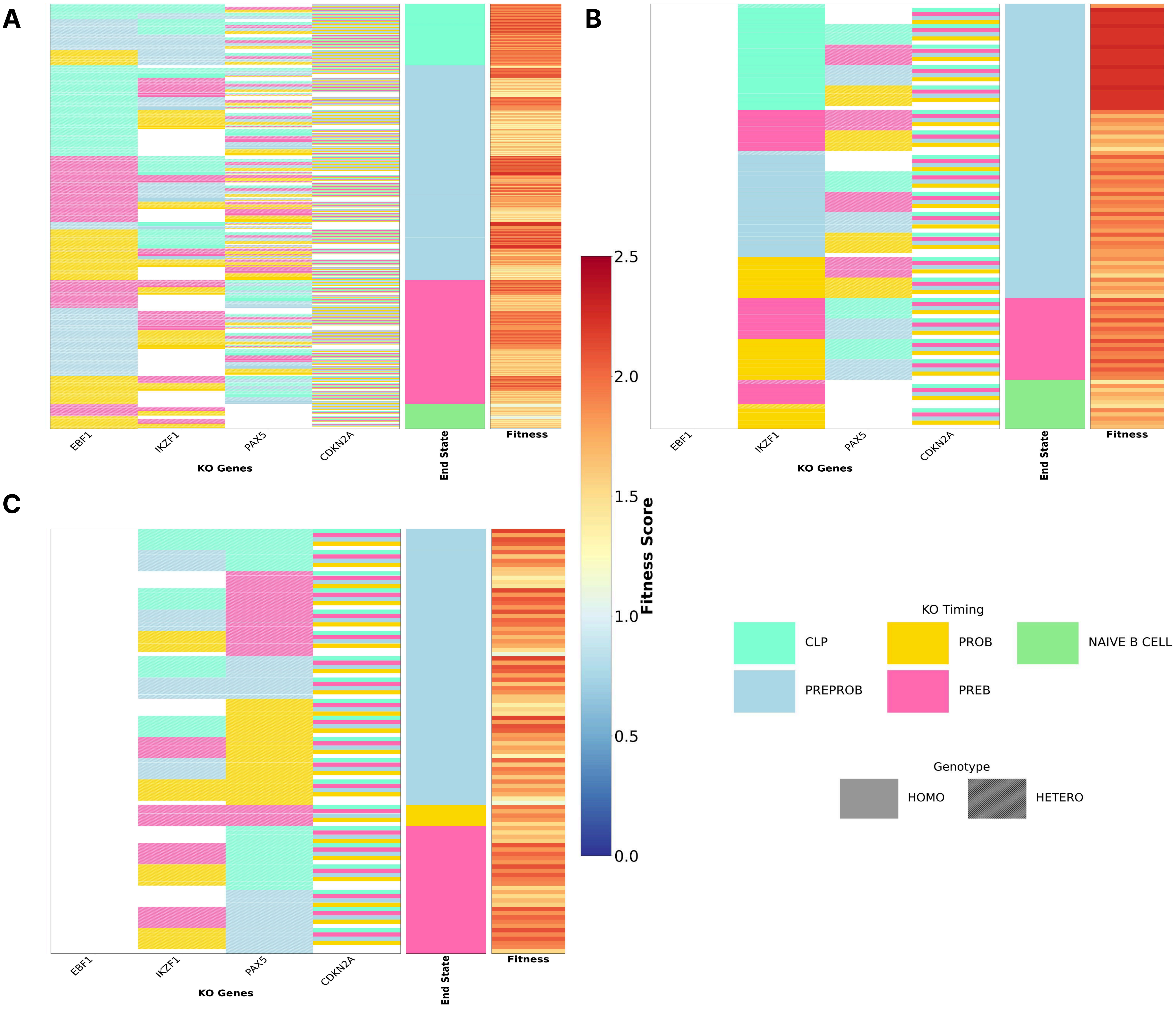

### S6Fig.pdf

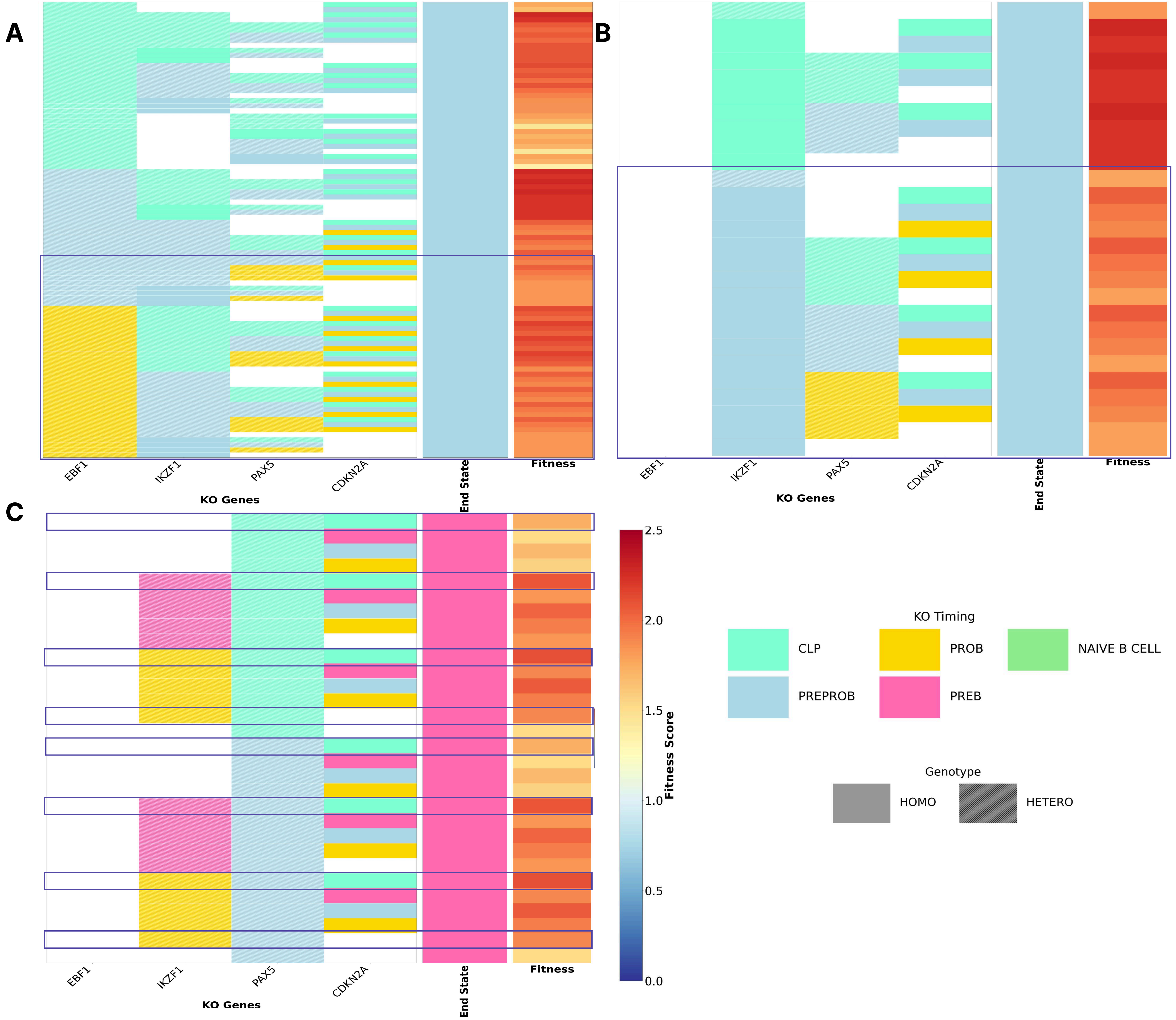

### S7Fig.pdf

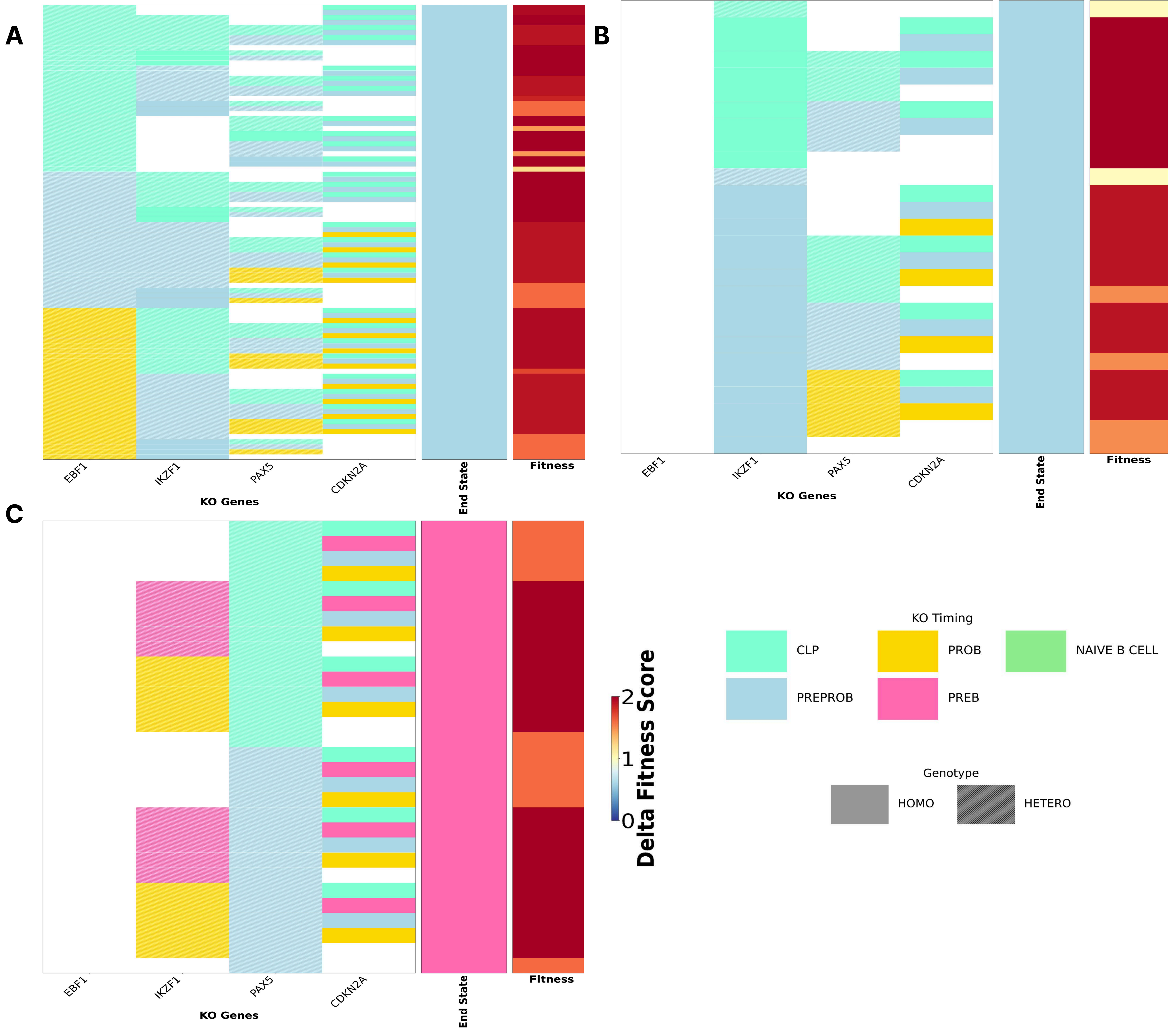

### S8Fig-bis.pdf

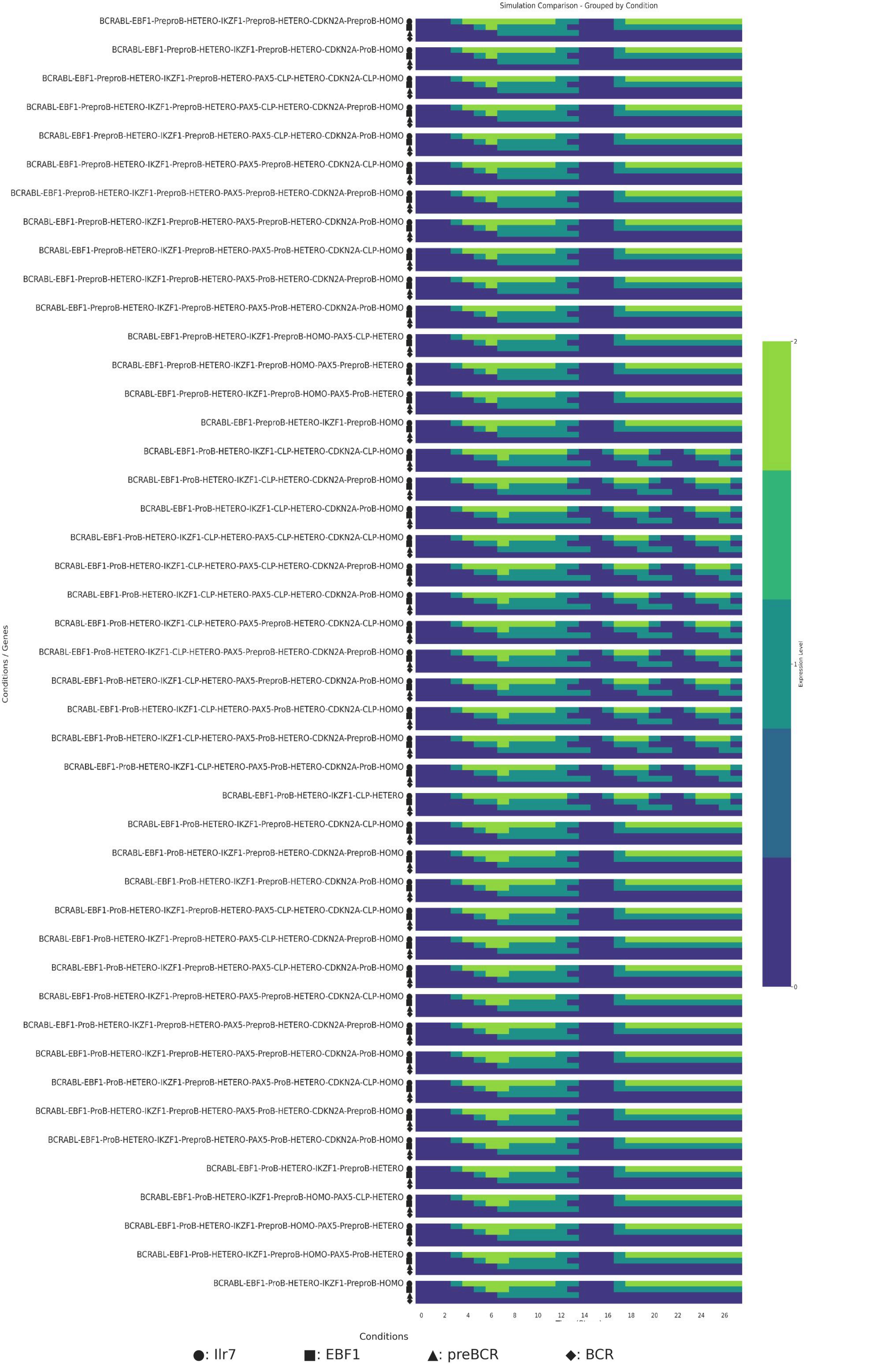

### S8Fig.pdf

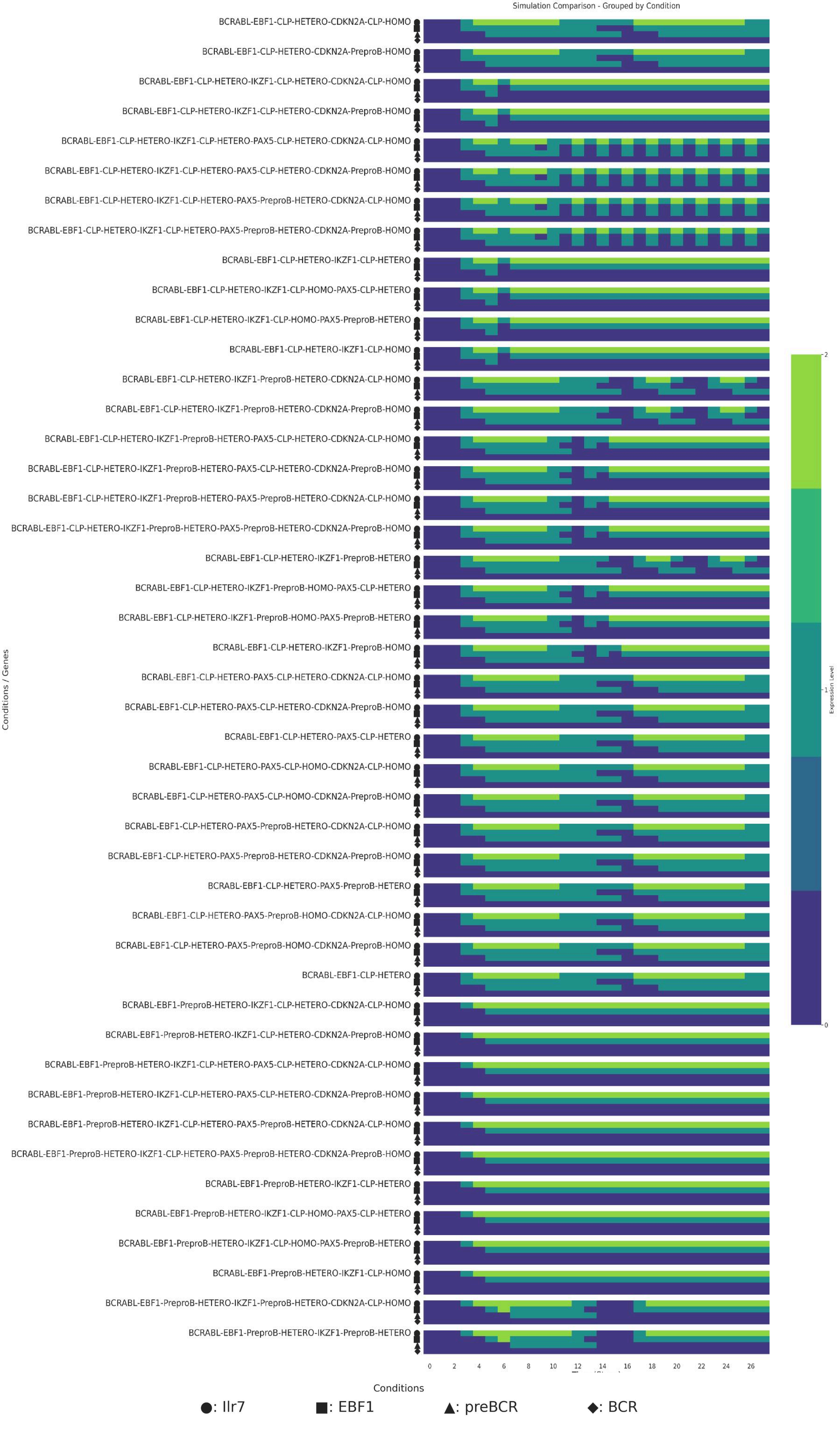

### S9Fig.pdf

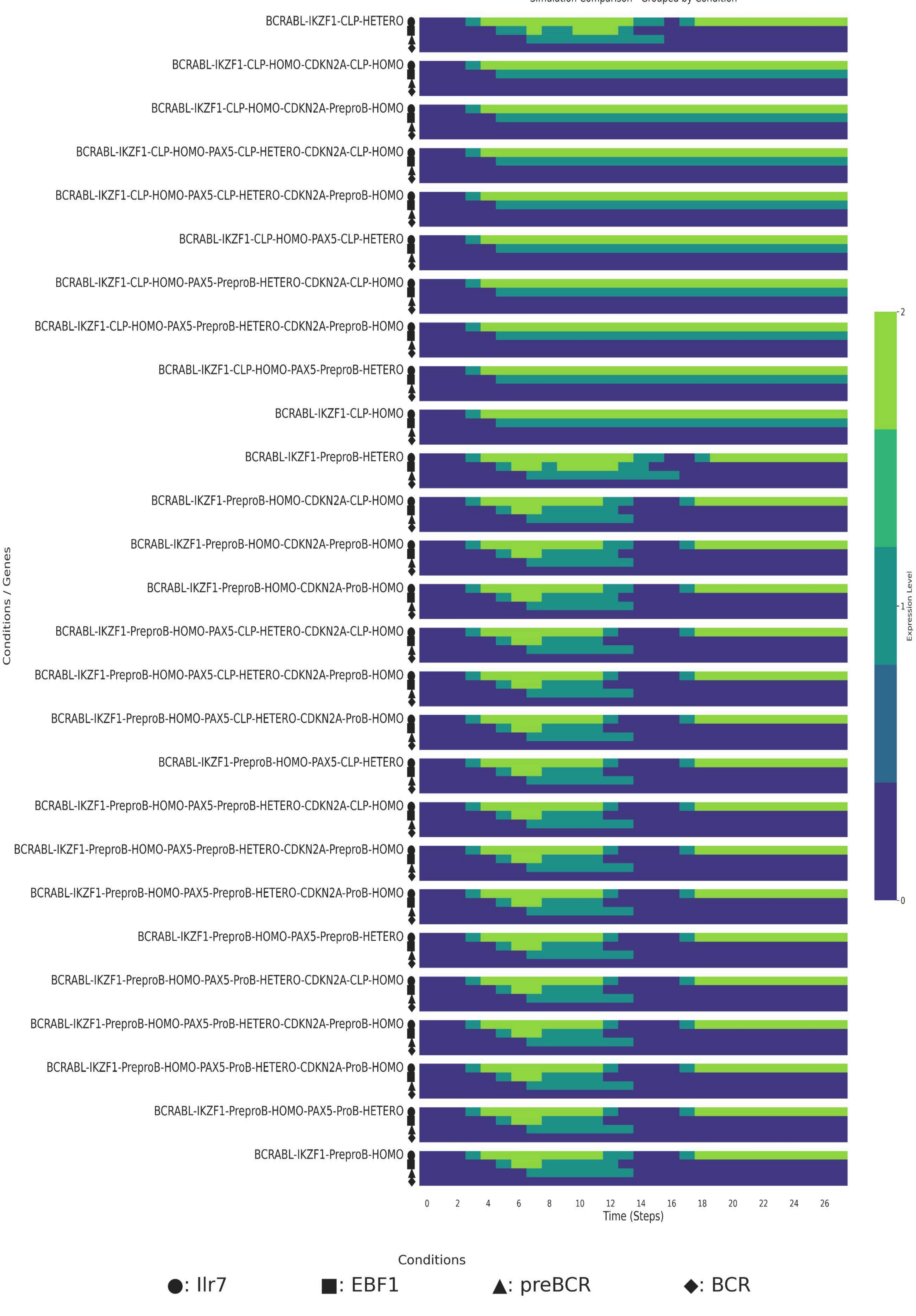

### S10Fig.pdf

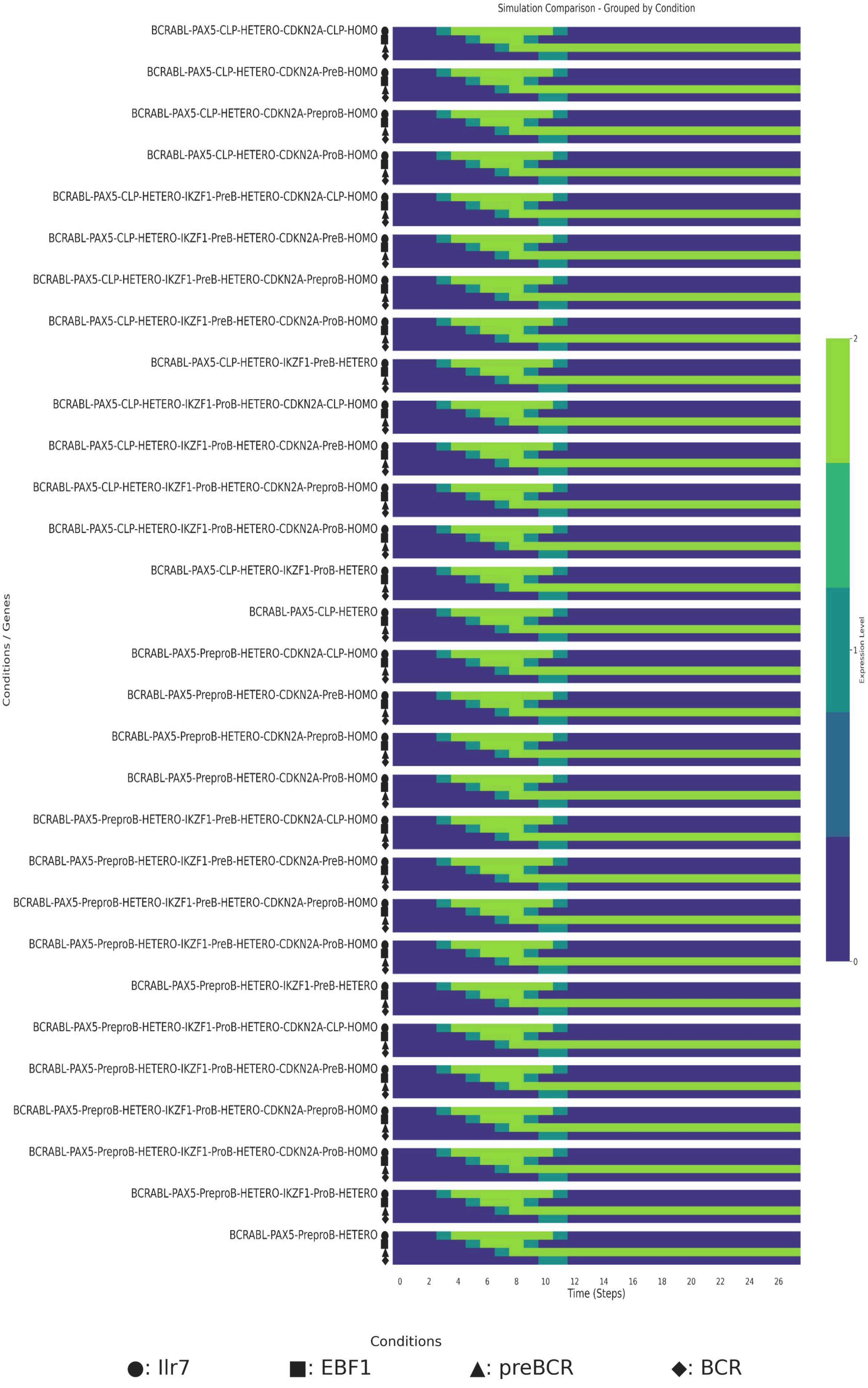

### S11Fig.pdf

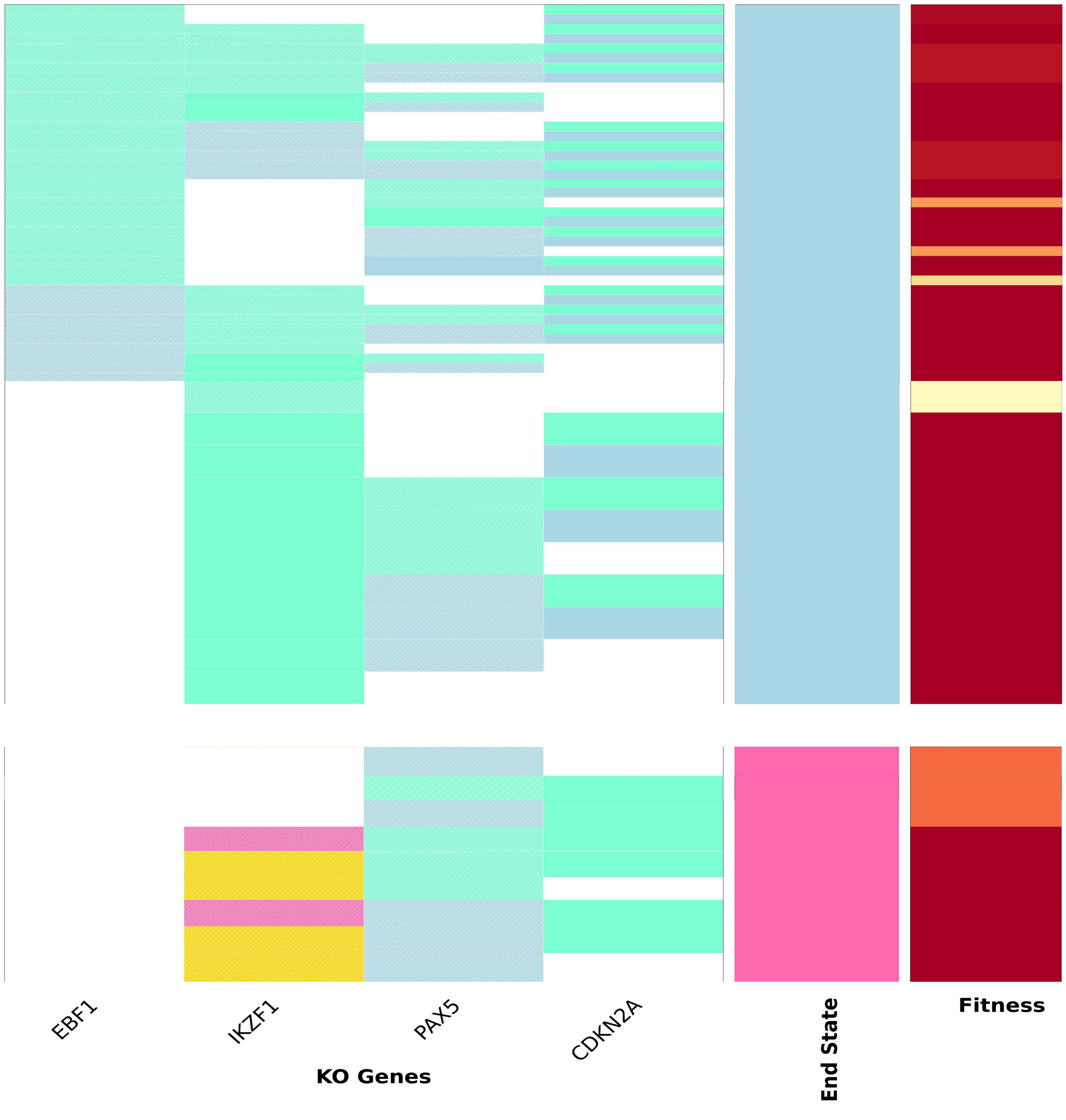
